# A distinct pool of Nav1.5 channels at the lateral membrane of murine ventricular cardiomyocytes

**DOI:** 10.1101/572180

**Authors:** Jean-Sébastien Rougier, Maria C. Essers, Ludovic Gillet, Sabrina Guichard, Stephan Sonntag, Doron Shmerling, Hugues Abriel

**Affiliations:** Institute of Biochemistry and Molecular Medicine, University of Bern, Bern CH-3012, Switzerland; Pain Center, Department of Anesthesiology, CHUV and the Department of Fundamental Neurosciences, University of Lausanne, Lausanne CH-1005, Switzerland; PolyGene AG, Rümlang, Switzerland

**Keywords:** voltage-gated sodium channel, lateral membrane of cardiomyocytes, α1-syntrophin, dystrophin, electrophysiology

## Abstract

**Background:** In cardiac ventricular muscle cells, the presence of voltage-gated sodium channels Na_v_1.5 at the lateral membrane depends in part on the interaction between the dystrophin-syntrophin complex and the Na_v_1.5 C-terminal PDZ-domain-binding sequence Ser-Ile-Val (SIV motif). α1-Syntrophin, a PDZ-domain adaptor protein, mediates the interaction between Na_v_1.5 and dystrophin at the lateral membrane of cardiac cells. Using the cell-attached patch-clamp approach on cardiomyocytes expressing Na_v_1.5 in which the SIV motif is deleted (ΔSIV), sodium current (I_Na_) recordings from the lateral membrane revealed an SIV-motif-independent I_Na_. Since immunostainings have suggested that Na_v_1.5 is expressed in transverse (T-) tubules, this remaining I_Na_ might be conducted by channels in the T-tubules. Of note, a recent study using heterologous expression systems showed that α1-syntrophin also interacts with the Na_v_1.5 N-terminus, which may explain the SIV-motif independent I_Na_ at the lateral membrane of cardiomyocytes.

**Aim:** To address the role of α1-syntrophin in regulating the I_Na_ at the lateral membrane of cardiac cells.

**Methods and results:** Patch-clamp experiments in cell-attached configuration were performed on the lateral membranes of wild-type, α1-syntrophin knock-down, and ΔSIV ventricular mouse cardiomyocytes. Compared to wild-type, a reduction of the lateral I_Na_ was observed in myocytes from α1-syntrophin knockdown hearts. However, similar to ΔSIV myocytes, a remaining I_Na_ was still recorded. In addition, cell-attached I_Na_ recordings from lateral membrane did not differ significantly between non-detubulated and detubulated ΔSIV cardiomyocytes. Lastly, we obtained evidence suggesting that cell-attached patch-clamp experiments on the lateral membrane cannot record currents conducted by channels in T-tubules such as calcium channels.

**Conclusion:** Altogether, these results suggest the presence of a sub-pool of sodium channels at the lateral membrane of cardiomyocytes that is independent of α1-syntrophin and the PDZ-binding motif of Na_v_ 1.5, located in membrane domains outside of T-tubules. The question of a T-tubular pool of Na_v_1.5 channels however remains open.

## Introduction

The sarcolemma of rod-shape adult ventricular cardiomyocytes can be divided into two principal regions. Firstly, the intercalated disc (ID) allows the electrical and physical coupling between adjacent cardiac cells *via* ID-ID connections. Secondly, the lateral membrane, including transverse (T-) tubules, contains focal adhesions crucial for the interaction between cardiomyocytes and the extracellular matrix [1-3]. The voltage-gated sodium channel Na_v_1.5 is expressed strongly in these membrane domains. Na_v_1.5 is the major isoform of the Na_v_1.x family in cardiomyocytes [4]. These channels are involved in the generation and propagation of the cardiac action potential (AP) [5, 6]. A few years ago, we and other groups investigating Na_v_1.5 started to unveil the complex localization of this channel in cardiac cells. We showed that at least two pools of Na_v_1.5 channels are present in murine ventricular cardiomyocytes based on their expression pattern and interaction with cytoplasmic proteins [7]. The first pool is located at the ID and plays a crucial role in the propagation of the electrical impulse. Based on results from heterologous expression systems and transfected cardiac cells, the regulation of this pool has been proposed to depend on SAP97 (synapse-associated protein 97), although *in vivo* experiments using cardiac-specific SAP97 knockdown mice did not support such observations [6, 8, 9]. Recently, Makara and colleagues have convincingly shown the crucial role of ankyrin-G in the regulation and targeting of Na_v_1.5 channel at the ID *in vivo* [10]. The second pool of Na_v_1.5 channels has been described outside the ID domain comprising the lateral membrane and the T-tubular system [5, 11, 12]. At the lateral membrane, the presence of Na_v_1.5 channels mainly depends on dystrophin and the PDZ-binding motif of Na_v_1.5 comprising its last three C-terminal amino acids Ser, Ile, and Val (SIV). The stability of this Na_v_1.5 pool has been shown to depend on the interaction between the dystrophin-syntrophin complex and the SIV motif [5, 11]. In this interaction, syntrophin, containing a PDZ domain, act as an adaptor protein between the SIV motif of Na_v_1.5 and dystrophin. Depending on the species, dystrophin can interact with different isoforms of syntrophin [13]. The syntrophin family comprises five isoforms of 60 kD each: α1, ß1, ß2, γ1, and γ2. α1-Syntrophin was the first syntrophin to be cloned from mouse heart due to its abundant expression [14]. The γ-isoforms are mainly expressed in the brain and skeletal muscle. α- and ß-isoforms are expressed in many tissues including the heart. In hamster, rat, and canine ventricular cardiomyocytes, dystrophin has been shown to interact preferentially with the α1-, β1-, and β2-isoforms of syntrophin [13]. In human and rabbit ventricles, mainly the α1-isoform interacts with dystrophin [13]. Using the cell-attached patch-clamp recording method on ventricular cardiomyocytes from genetically modified mice expressing Na_v_1.5 lacking the SIV motif (ΔSIV mouse model), a ∼60% reduction, but not an abolition, of I_Na_ has been observed at the lateral membrane [11]. In addition, the biophysical properties of the remaining I_Na_ at the lateral membrane of ΔSIV myocytes were similar to the wild-type cells from littermate mice [11]. One hypothesis proposed by Shy *et al.* to explain the presence of this remaining current was the presence of Na_v_1.5 channels at the T-tubules. However, the presence of Na_v_1.5 in T-tubular system is still under debate. Although immunofluorescent stainings for Na_v_1.5 have shown a T-tubular-like pattern in ventricular cardiomyocytes, other experiments suggest that other Na_v_1.x isoforms might be the main isoforms in this structure [11, 15, 16]. Interestingly, Matamoros and colleagues,, have shown in heterologous expression systems that α1-syntrophin can not only interact with the SIV motif but also with the sequence comprising three N-terminal amino acids Ser, Leu, Ala (SLA) of Na_v_1.5 channels [17]. Based on this observation, the remaining sodium current observed on the lateral membrane of ΔSIV mice might also depend on α1-syntrophin interacting with Na_v_1.5 through the N-terminal SLA motif.

Taking these published results together, we might explain the remaining I_Na_ outside the ID in ΔSIV ventricular cardiomyocytes by the following working hypothesis explained by 1) Na_v_1.5 channels interact with the dystrophin-syntrophin complex *via* its N-terminal SLA motif; and/or 2) Na_v_1.5 channels localize within a specific area of the lateral membrane independent of the dystrophin-syntrophin complex; and/or 3) Na_v_1.5 channels are present inside the T-tubular system. In this work, we aim to test these hypotheses.

## Methods

### Animals

All experiments involving animals were performed according to the Swiss Federal Animal Protection Law and had been approved by the Cantonal Veterinary Administration, Bern. This investigation conforms to the Guide for the Care and Use of Laboratory Animals, published by the US National Institutes of Health (NIH publication no. 85-23, revised 1996).

### Generation of constitutive cardiac-specific α1-syntrophin knockdown mice

To study the function of α1-syntrophin (Sntα1), a conditional mouse model was generated that allows tissue or time-point specific inactivation of the *Snta1* gene by the use of the Cre-/loxP-system. The design of the mouse model is based on a modified *Snta1* locus with two loxP sites flanking part of *Snta1* exon1. One loxP site is inserted in the 5′ UTR in exon1 whereas the second loxP site is located in intron 1. Upon breeding with a heart-specific Cre-transgenic mouse line (i.e., MHC-Cre), the coding region on exon 1 of *Snta1* is deleted, restricted to the cardiac muscle that is studied. For the introduction of loxP sites into the *Snta1* locus, a targeting vector (pSnta1 TV) was generated that contained homology regions of 2.2 kb (short homology region) and 5.6 kb (long homology region). The 3′ loxP site was inserted together with an FRT-flanked neomycin cassette into a KpnI restriction site 269 bp downstream of *Snta1* exon 1. The 5′ loxP site was inserted 50 bp downstream of the start of exon 1 and 78 bp upstream of the ATG start codon in the 5′ untranslated region (5′ UTR). In total a region of 639 bp was flanked by loxP sites, containing the coding part of *Snta1* exon 1. After transfection of 20 mg of the targeting vector into 10^7^ 129Ola derived embryonic stem cells and selection with 200 mg/ml G418 for 10 days, 100 clones were isolated and screened for insertion into the *Snta1* locus via Southern blot analysis using PstI digested genomic DNA and a 5′ external probe (Fig. S1A). Eight positive clones were identified and further analyzed by Southern blot using PstI digested genomic DNA and a 3′ external probe (Fig. S1B). Two clones with a correct karyotype were injected into C57BL/6 derived blastocysts. Male mice with a high degree of coat color chimerism were mated to Flp-deleter mice (C57Bl/6 background) to generate heterozygous *Snta1*-flox mice. For genotyping, two primers flanking the 5′ loxP site were used (Fig. S1A). Primers E072.32 (5′-ACGAGCTACGGTCCAATC-3′) and E072.33 (5′-ATCTTCGCCTCGAAGTCC-3′) detect a 391 bp fragment in the wild-type allele whereas the insertion of the loxP site in the *Snta1*-flox allele results in a slightly larger fragment of 437 bp.

### Cell line preparations

Human embryonic kidney (HEK-293) cells were cultured in DMEM (Gibco, Basel, Switzerland) supplemented with 10% FBS, 0.5% penicillin, and streptomycin (10,000 U/mL) at 37°C in a 5% CO2 incubator.

### Transfections

For electrophysiological studies, T25 cm2 flasks of HEK-293 cells were transiently co-transfected using Jet-Pei^®^ reagent (Polyplus Transfection, Illkirch, France) with 10 µg of human Na_v_1.4 cDNA. All transfections included 0.3 µg of cDNA encoding CD8. For patch clamp experiments, anti-CD8 beads (Dynal^®^, Oslo, Norway) were used. Only cells decorated with anti-CD8 beads were analyzed.

### Isolation of mouse ventricular myocytes

Single cardiomyocytes were isolated according to a modified procedure of established enzymatic methods [5]. Briefly, mice were euthanized by cervical dislocation. Hearts were rapidly excised, cannulated and mounted on a Langendorff column for retrograde perfusion at 37°C. Hearts were rinsed free of blood with a nominally Ca^2+^-free solution containing (in mM): 135 NaCl, 4 KCl, 1.2 MgCl_2_, 1.2 NaH_2_PO_4_, 10 HEPES, 11 glucose, pH 7.4 (NaOH adjusted), and subsequently digested by a solution supplemented with 50 µM Ca^2+^ and collagenase type II (1 mg/mL, Worthington, Allschwil, Switzerland) for 15 minutes. Following digestion, the atria were removed and the ventricles transferred to nominally Ca^2+^-free solution, where they were minced into small pieces. Single cardiac myocytes were liberated by gentle trituration of the digested ventricular tissue and filtered through a 100 µm nylon mesh. Ventricular mouse cardiomyocytes were used after an extracellular calcium increase procedure to avoid calcium overload when applying extracellular solutions in electrophysiology assays.

### Detubulation

Isolated cardiomyocytes were detubulated *via* osmotic shock using formamide solution [18]. Briefly, cardiac cells were suspended in 2 mL of 1.5 M formamide solution for 15 minutes at room temperature (RT) then washed twice with an extracellular solution before use. For morphological experiments, 1 µM di-8 ANEPPS, diluted in dimethyl sulfoxide containing pluronic F-127, was applied to living cardiomyocytes for 15 minutes, and imaged on a confocal microscope (LSM710, Zeiss, Germany).

### Whole-cell electrophysiology

Cardiac action potentials (AP) and ionic currents (I_Na_, calcium current (I_Ca_), transient outward potassium current (I_to_), and the inward rectifier potassium current (I_K1_)) were recorded in the whole-cell configuration at RT (22-23°C) using a VE-2 amplifier. Borosilicate glass pipettes were pulled to a series resistance of ∼2 MΩ pClamp software, version 8 (Axon Instruments, Union City, CA, USA) was used for recordings. Data were analyzed using pClamp software, version 8 (Axon Instruments) and OriginPro, version 7.5 (OriginLab Corp., Northampton, MA, USA). Current densities (pA/ pF) were calculated by dividing the peak current by the cell capacitance. To determine the voltage dependence of steady-state activation and inactivation curves, the data from individual cells were fitted with the Boltzmann equation, y(V_m_) = 1/(1+exp[(V_m_-V_1/2_)/K]), where y is the normalized current or conductance, V_m_ is the membrane potential, V_1/2_ is the voltage at which half of the channels are activated or inactivated, and K is the slope factor.

For cardiac AP recordings, cardiomyocytes were bathed in a solution containing (in mM) NaCl 140, KCl 5.4, CaCl_2_ 1.8, MgCl_2_ 1.2, HEPES 10, and glucose 5 (pH was adjusted to 7.4 with NaOH). Cardiomyocytes were initially voltage-clamped (holding potential −80 mV) and dialyzed with an internal solution containing (in mM) KCl 120, CaCl_2_ 1.5, MgCl_2_ 5.5, Na_2_ATP 5, K_2_-EGTA 5, and HEPES 10 (pH was adjusted to 7.2 with KOH). APs were elicited at 0.5 Hz with rectangular pulses (5 ms at 125% threshold) in current-clamp mode. Elicited APs were allowed to stabilize before one or more sequences of ∼1 minute each were acquired from each cell. AP recordings were digitized at a sampling frequency of 20 kHz. Electrophysiological data were analyzed off-line, where the resting membrane potential, amplitude, maximal upstroke velocity (dV/dt)_max_ and durations at 30, 50 and 90% repolarization were averaged from each sequence of APs.

I_Na_ in α1-syntrophin knockdown (*Snta1* fl/fl^+^) or α1-syntrophin wild-type littermate (*Snta1* fl/fl^-^) cardiomyocytes were carried out using an internal solution containing (in mM) CsCl 60, Cs-aspartate 70, CaCl_2_ 1, MgCl_2_ 1, HEPES 10, EGTA 11, and Na_2_ATP 5 (pH was adjusted to 7.2 with CsOH). Cardiomyocytes were bathed in a solution containing (in mM) NaCl 5, NMDG-Cl 125, CaCl_2_ 2, MgCl_2_ 1.2, CsCl 5, HEPES 10, and glucose 5 (pH was adjusted to 7.4 with CsOH). Nifedipine (10 µM) and cobalt chloride (CoCl_2_ ) (10 µM) were added to the extracellular solution in order to inhibit calcium currents.

To assess the functional consequences of detubulation, I_Ca_ was recorded using the internal pipette solution composed of (in mM) 60 CsCl, 70 Cs-aspartate, 1 MgCl_2_, 10 HEPES, 11 EGTA and 5 Mg-ATP (pH was adjusted to 7.2 with CsOH). The external solution contained (in mM) 5 NaCl, 125 NMDG-Cl, 5.6 CsCl, 5 BaCl_2_, 1 MgCl_2_, 10 HEPES and 5 D-glucose (pH was adjusted to 7.4 with CsOH).

I_to_ and I_K1_ recordings were performed using an internal solution containing (in mM): KCl 130, Mg-ATP 4, NaCl 12, MgCl_2_ 1, CaCl_2_ 1, EGTA 10, HEPES 10 (pH was adjusted to 7.2 with KOH). Myocytes were bathed with a solution containing (in mM): NaCl 140, KCl 5.4, CaCl_2_ 1.8, MgCl_2_ 1, HEPES 10, and Glucose 5 (pH was adjusted to 7.4 with NaOH). Tetrodotoxin (TTX) sensitive sodium channels (Na_v_1.1, Na_v_1.2, Na_v_1.3, Na_v_1.4, Na_v_ 1.6, and Na_v_ 1.7) were blocked with 50 µM of TTX. Calcium channels were blocked with 3 mM CoCl_2_. I_K1_ barium-sensitive current was calculated by subtracting potassium current recorded after perfusion of an extracellular solution containing 100 µM barium chloride (BaCl_2_) from potassium current recorded before application of BaCl. To record Ito, 100 µM BaCl_2_ was added to the extracellular solution to block I_K1_ current.

### Cell-attached electrophysiology

I_Na_ macro recordings were recorded in the cell-attached configuration either on the lateral membrane of cardiomyocytes or in HEK-293 cells transiently expressing Na_v_1.4. All cell-attached recordings were performed at RT (22-23°C), using a low-noise Axopatch 200B amplifier (Molecular Devices, Sunnyvale, CA, USA). Borosilicate glass pipettes were pulled to a series resistance of ∼1.5 MΩ (diameter of opening ∼20 µm^2^). The internal pipette (intrapipette) solution was composed of (in mM) 148 NaCl, 1.8 CaCl_2_, 1 MgCl_2_, 5 D-glucose, and 10 HEPES (pH was adjusted to 7.4 with NaOH). The external solution, to depolarize the cell membrane ∼0 mV, contained (in mM) 125 KCl, 30 KOH, 10 EGTA, 2 CaCl_2_, and 5 HEPES (pH was adjusted to 7.15 with KOH). A 4-pole Bessel low-pass filter at 2 kHz, was applied during the recording and a sampling rate of 20 kHz was used for the acquisition of the raw traces. P over 4 protocol was applied. pClamp software, version 8 (Axon Instruments, Union City, CA, USA) was used for recordings. Data were analyzed using pClamp software, version 8 (Axon Instruments) and OriginPro, version 7.5 (OriginLab Corp., Northampton, MA, USA). To determine the voltage dependence of steady-state activation and inactivation curves, the data from individual cells were fitted with the Boltzmann equation, y(V_m_) =1/(1+exp[(V_m_-V_1/2_)/K]), where y is the normalized current or conductance, V_m_ is the membrane potential, V_1/2_ is the voltage at which half of the channels are activated or inactivated, and K is the slope factor.

Single channel barium recordings have been performed in cell-attached configuration. Average pipette size was 1.5 MΩ. Currents were measured at RT (22–23°C) using a low-noise Axopatch 200B amplifier (Molecular Devices, Sunnyvale, CA, USA). The internal pipette solution was composed of (in mM) 110 BaCl, 10 TEA-Cl, and 5 HEPES (pH was adjusted to 7.4 with NMDG-OH). The external solution, to depolarize the cell membrane ∼0 mV, contained (in mM) 125 KCl, 30 KOH, 10 EGTA, 2 CaCl_2_, and 5 HEPES (pH was adjusted to 7.15 with KOH). A 4-pole Bessel low-pass filter at 2 kHz, applied during the recording and a sampling rate of 20 kHz was used for the acquisition of the raw traces. For each cell, a single depolarizing protocol at −10 mV from a holding potential at −120 mV was applied at least 150 times. Post-acquisition, data were filtered using a low pass Gaussian filter and analyzed using pClamp software, version 9.2 (Axon Instruments, Union City, CA, USA).

### Proximity ligation assay

Duolink^®^ (Olink Bioscience, Uppsala, Sweden) is a proximity ligation assay (PLA) that enables the visualization of protein-protein interactions allowing the *in situ* detection of epitopes that are near each other (<40 nm apart). Two primary antibodies raised in different species recognize the target antigen or antigens of interest. Species-specific secondary antibodies, called PLA probes, each with a unique short DNA strand attached to it, bind to the primary antibodies. When PLA probes are nearby, the DNA strands can interact through subsequent addition of two other circle-forming DNA oligonucleotides. After joining of the two oligonucleotides by enzymatic ligation, they are amplified *via* rolling circle amplification using a polymerase. After the amplification reaction, several-hundredfold replication of the DNA circle has occurred, and labeled complementary oligonucleotide probes highlight the product. The resulting high concentration of fluorescence in each single-molecule amplification product is easily visible as a distinct bright dot when viewed with a fluorescence microscope. Experiments were performed according to the manufacturer’s recommendations. Briefly, isolated mouse ventricular cardiac cells were fixed in 4% paraformaldehyde for 10 minutes and incubated twice for 7 minutes in PBS with 0.02% Triton X-100 at RT. Sub-sequently, cells were incubated 15 minutes in blocking solution containing 1% bovine serum albumin (BSA). Cardiomyocytes from α1-syntrophin knockdown or α1-syntrophin wild-type littermate were then incubated with commercially available mouse monoclonal anti-pan-syntrophin and polyclonal anti-Na_v_1.5 primary antibodies as specified below at a dilution of 1/100 and 1/200, respectively. Subsequent incubation steps were performed at 37°C and the appropriate washing buffers were used after each incubation, as specified in the protocol. After washing 3 times in PBS to remove the primary antibody, Duolink^®^ secondary antibodies conjugated to PLA probes were added to the tissue for a 1-hour incubation. These PLA probes consist of oligonucleotides that were subsequently joined together in a circle after the addition of a ligation solution (containing ligase enzyme) and incubation for 30 minutes. Next, the rolling circle amplification of this circular template was achieved by the addition of an amplification solution, containing polymerase and fluorescently labeled oligonucleotides, which hybridized to the circular template during a 100-minute incubation. Samples were then mounted with Duolink^®^ *in situ* mounting medium with DAPI (1 µl/200 µl PBS) and viewed with a confocal microscope (LSM710, Zeiss, Germany).

### Protein extraction and western blots

Whole hearts were extracted from mice, and the atria were removed before homogenization of the ventricles in 1.5 mL lysis buffer (50 mM HEPES, 150 mM NaCl, 1 mM EGTA, 10% glycerol, 1.5 mM MgCl_2_, and Complete^®^ protease inhibitor cocktail from Roche, Basel, Switzerland) using a Polytron. Triton X-100 was then added to make a final concentration of 1% (final volume 3 mL) before lysis of the samples on a rotating wheel for 1 hour at 4°C. Soluble fractions of the mouse heart were obtained by subsequent centrifugation for 15 minutes at 13,000 rpm at 4°C. To determine the correct volume of sample needed to load equivalent amounts of protein on a gel, the protein concentration of each of the lysate samples was measured in triplicate by Bradford assay and intrapolated by a BSA standard curve. Samples were denatured at 95°C for 5 minutes before loading them on a gel. To analyze and compare protein content in whole mouse hearts, 25 µg or 50 µg of ventricular mouse heart lysate samples were loaded on a 1.0 mm-thick 10% Tris-acetate gel for high molecular weight proteins (Invitrogen). Gels were run at 150 V for 1 hour and then subsequently transferred to nitrocellulose membranes using the Biorad Turbo Transfer System (Biorad, Hercules, CA, USA). All membranes were stained with Ponceau as a qualitative check for equivalent loading of total protein. Membranes were then rinsed twice with PBS before using the SNAP i.d. system (Millipore, Zug, Switzerland) for western blotting. 1% BSA was used as a blocking solution and incubated with membranes for 10 minutes before incubation with the primary antibodies for 10 minutes. Membranes were subsequently washed 4 times in PBS + 0.1 % Tween before incubating with fluorescent secondary antibodies for 10 minutes. After 4 more washes with PBS + 0.1% Tween and 3 washes in PBS, membranes were scanned with the Odyssey^®^ Infrared Imaging System (LI-COR Biosciences, Bad Homberg, Germany) for detection of fluorescent protein. Subsequent quantitative analysis of protein content was achieved by measuring and comparing band densities (equivalent to fluorescence intensities of the bands) using Odyssey software version 3.0.21.

### Immunocytochemistry

Following cardiomyocyte isolation, cells from α1-syntrophin knockdown and α1-syntrophin wild-type littermate hearts were fixed with cold methanol at −20°C for 10 minutes. Cells were subsequently washed twice in PBS before permeabilization with PBS containing 0.02% Triton X-100 for 14 minutes. Then, the cardiomyocytes were blocked for 15 minutes at RT with 10% goat serum in TBS containing 2.3 % of fab fragment donkey anti-mouse IgG (H+L) (Jackson Immuno Research, Baltimore, USA) at 1.3 mg/mL. Cardiomyocytes were then incubated overnight with primary antibodies diluted in 3% goat serum in TBS containing 0.02% Triton X-100. Afterwards, cells were washed 4 times with 3% goat serum in TBS for 5 minutes each, and then incubated with secondary antibodies for 2 hours at RT. Cells were then washed 4 times in PBS (5 minutes each). Finally, DAPI solution (1 µl/200 µl PBS) was applied on cardiac cells for 20 minutes at RT. Before examination with the confocal microscope (LSM710, Zeiss, Germany), cardiomyocytes were embedded in FluorSave^TM^ Reagent (Calbiochem). Total signal intensity was calculated using Zen 2.1 software version 11 from Carl Zeiss (Germany).

### Antibodies

For western blots, the antibody against pan-syntrophin (MAI-745, Affinity BioReagents, Golden, CO, USA) was used at a dilution of 1/200. A custom rabbit polyclonal antibody against α1-syntrophin (Pineda Antibody Service, Berlin) was used at a dilution of 1/1000. The antibodies against Cre (69050; Novagen, EMD Millipore-MERK, Schaffhausen, Switzerland) was used at a dilution of 1/500 and the antibody against calnexin (C4731; Sigma-Aldrich, Saint-Louis, MO, USA) was used at 1/1000. For Duolink^®^ experiments, the antibody against syntrophin (pan-syntrophin, MAI-745, Affinity BioReagents, Golden, CO, USA) was used at a dilution of 1/100 and custom rabbit polyclonal antibody against Na_v_1.5 (Pineda Antibody Service, Berlin) was used at a dilution of 1/200. For stainings, the anti-pan-syntrophin antibody (MAI-745, Affinity BioReagents, Golden, CO, USA) was used at a dilution of 1/200. The antibody against Cre (69050; Novagen, EMD Millipore-MERK, Schaffhausen, Switzerland) was used at a dilution of 1/1000.

### Drugs

Tetrodotoxin (TTX) was purchased from Alomone Labs (Jerusalem, Israel) and dissolved in intrapipette solution. The µ-conotoxin CnIIIC was purchased from Smartox (Saint Martin d’Heres, France) and dissolved in water.

### Statistical analyses

Data are represented as means ± S.E.M. Statistical analyses were performed using Prism7 GraphPad(tm) software. A Mann-Whitney U two-tailed test was used to compare two groups. *p* < 0.05 was considered significant.

## Results

### Characterization of a cardiac-specific α1-syntrophin knockdown mouse strain

The remaining I_Na_ at the lateral membrane of Na_v_1.5 ΔSIV mouse cardiomyocytes [11] raised the question which molecular determinants are involved in this lateral membrane pool of Na_v_1.5 channels. Although the Na 1.5 SIV motif mediates the interaction with α1-syntrophin of the dystrophin-syntrophin complex, Na_v_1.5 contains a similar motif (Ser-Leu-Ala) in its N-terminal domain, which also interacts with α1-syntrophin [17]. Motivated by these observations, we generated a cardiac-specific α1-syntrophin knockdown mouse strain (described in Supplemental Fig.1) and investigated the consequences of α1-syntrophin knockdown on the I_Na_ of ventricular cardiomyocytes. Using either pan-syntrophin (targetting all members of the syntrophin family) or α1-syntrophin antibodies, western blots confirmed the cardiac-specific reduction of the α1-syntrophin expression in *Snta1* fl/fl^+^ cardiac tissues compared to the control *Snta1* fl/fl^-^ (Fig. 1A and 1B). In addition, immunostaining experiments using the pan-syntrophin antibody showed a significant decrease of syntrophin expression in isolated ventricular cardiomyocytes *Snta1* fl/fl^+^ compared to *Snta1* fl/fl^-^ cardiac cells (Fig. 2A and 2B). Finally, proximity ligation assay experiments showed a substantial decrease of the interaction (based on the absence of red dots) between Na_v_ 1.5 and α1-syntrophin in *Snta1* fl/fl^+^ cardiomyocytes compared to the control, suggesting that α1-syntrophin and Na_v_ 1.5 no longer interact in α1-syntrophin knockdown cells (Fig. 2C). In parallel to the molecular characterization, functional investigations were performed. Whole-cellI_Na_ was recorded in freshly isolated ventricular cardiomyocytes. As shown in figures 3A and 3B, comparing knockdown cardiomyocytes to the wild-type cells, a significant reduction of the I_Na_ was observed (*Snta1* fl/ fl^-^ MaxI _Na_: 33 ± 2 pA/pF, n = 16; *Snta1* fl/fl^+^ MaxI_Na_: 22 ± 2 pA/pF, n = 13, p < 0.05), without alteration of I_Na_ biophysical properties (Fig. 3A and 3B and supplemental Fig. 2). Cardiac AP recordings moreover did not show any alteration of neither the resting membrane potential nor AP duration (Table 1, Fig. 4A and 4B). Only the maximal upstroke velocity was significantly decreased in α1-syntrophin knockdown cells compared to controls (Table 1 and Fig. 4B). Using heterologous expression systems, Matamoros *et al*. have also demonstrated that α1-syntrophin interacts with Kv4.2/4.3 and Kir2.1 potassium channels, which conduct the Ito and I_K1_ currents, respectively [17]. Although the decrease of the upstroke velocity of the AP suggests that only I_Na_ is altered in α1-syntrophin knockdown conditions, we recorded the Ito and I_K1_ currents in the absence of α1-syntrophin. As shown in figures 5 and 6, no alteration was observed in α1-syntrophin knockdown cells compared to the control for both currents (*Snta1* fl/fl^-^ MaxI_to_: 51 ± 6 pA/pF, n = 12; *Snta1* fl/fl^+^ MaxIto: 50 ± 6 pA/pF, n = 13, p > 0.05) (Fig. 5A and 5B), and (*Snta1* fl/fl^-^ MaxI_k1_: 6.4 ± 0.6 pA/pF, n=12; *Snta1* fl/fl^+^ MaxI_K1_: 7.0 ± 0.6 pA/pF, n = 15, p > 0.05) (Fig. 6A and 6B) suggesting that mainly the Na_v_1.5-mediated current is down-regulated in this α1-syntrophin knockdown model.

**Table 1.** Cardiac action potential parameters. Resting membrane potential (RMP), action potential duration (APD) and maximum upstroke velocity (dV/dt) calculated from ventricular cardiac action potentials of wild type (*Snta1* fl/fl^-^) and syntrophin knockdown (*Snta1* fl/fl^+^) mice. Number of cells in parenthesis. *; *p* < 0.05.

|  | *Snta1* fl/fl<sup>-</sup> | *Snta1* fl/fl<sup>+</sup> |
| --- | --- | --- |
| RMP (mV) | -90±1 (n=21) | -89±1 (n=22) |
| APD<sub>30</sub> (ms) | 3.6±0.5 (n=21) | 4.2±0.8 (n=22) |
| APD<sub>50</sub> (ms) | 7.6±1.1 (n=21) | 9.1±1.7 (n=22) |
| APD<sub>90</sub> (ms) | 35.5±2.6 (n=21) | 33.2±3.3 (n=22) |
| dV/dt (mV/ms) | 218±7 (n=21) | 199±5 (n=22) * |

**Figure 1.**
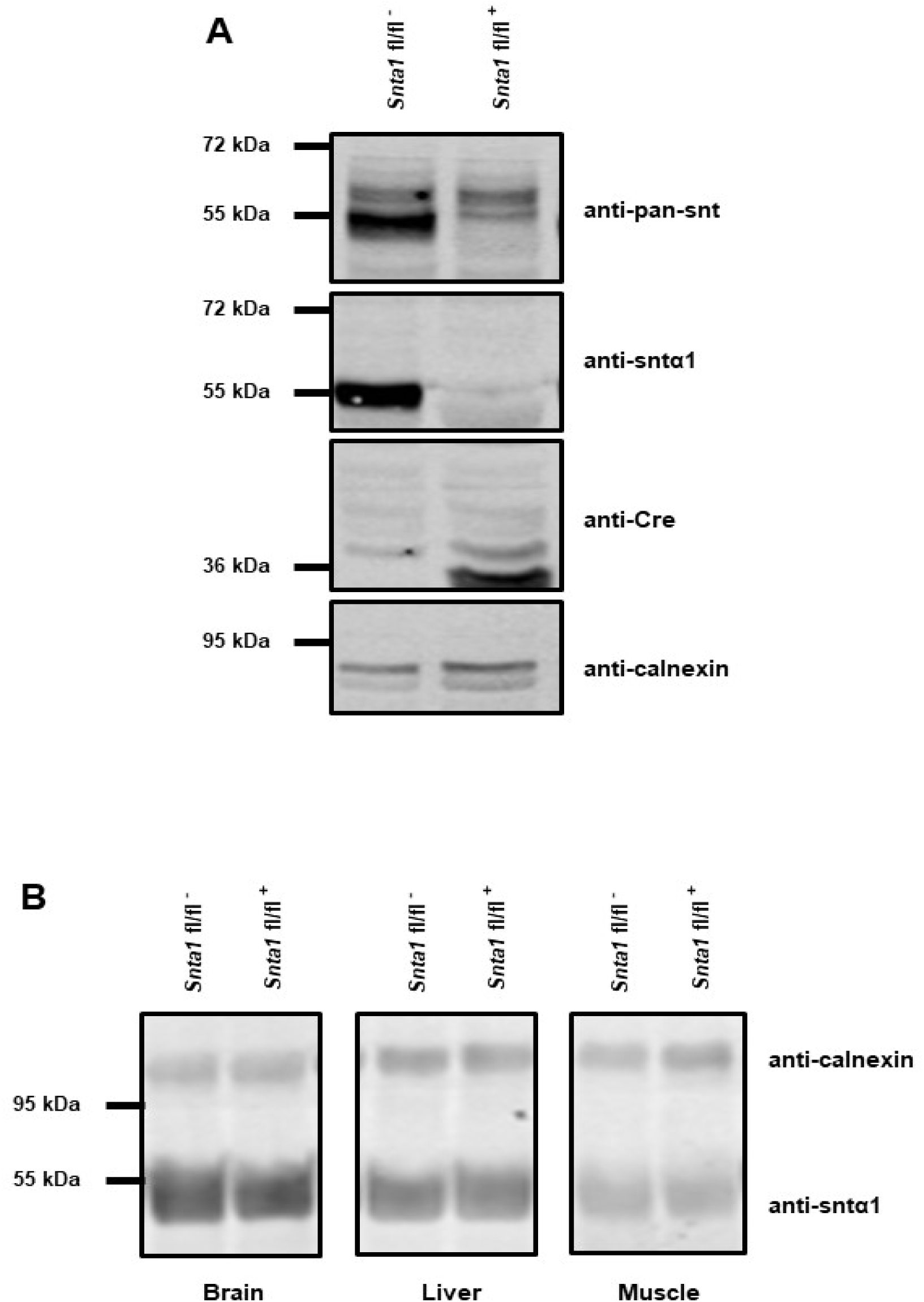
α1-syntrophin knockdown characterization. A, Western blots of whole hearts from wild-type (*Snta1* fl/fl^-^) and α1-syntrophin cardiac specific knockdown (*Snta1* fl/fl^+^) mice show a significant decrease of the α1-syntrophin protein (anti-Snta1 antibody) and the absence of compensatory effects *via* other syntrophin isoforms (anti-pan-syntrophin (Snt) antibody). B, Western blots from brain, liver and muscle from wild-type (*Snta1* fl/fl^-^) and α1-syntrophin cardiac specific knockdown (*Snta1* fl/fl^+^) mice show no effect on α1-syntrophin protein in other organs than the heart.

**Figure 2.**
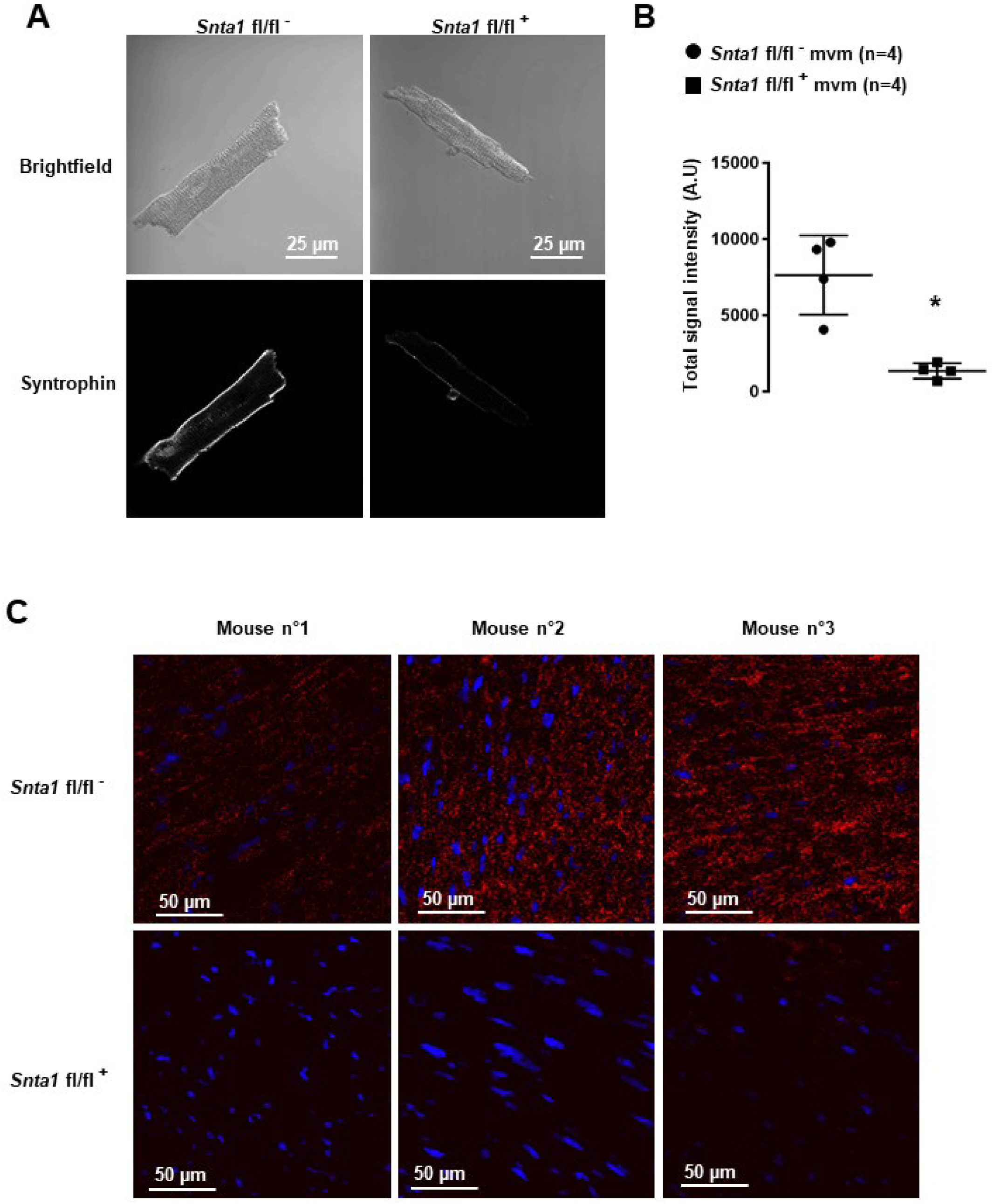
Syntrophin staining. A, Immunostainings, using a pan-syntrophin antibody, of syntrophin in wild-type (*Snta1* fl/fl^-^) and cardiac-specific α1-syntrophin knockdown murine cardiomyocytes (mvm) (*Snta1* fl/fl^+^). B, Intensity quantification of syntrophin signal of four cells confirming the significant decrease of syntrophin in knockdown mice. (*; *p* < 0.05). The number of cells is indicated in parentheses. C, Duolink® experiments showing a significant decrease in interaction (red dots) between Na _v_1.5 and syntrophin in α1-syntrophin cardiac specific knockdown cardiomyocytes (*Snta1* fl/fl^+^) compared to wild type (*Snta1* fl/fl^-^).

**Figure 3.**
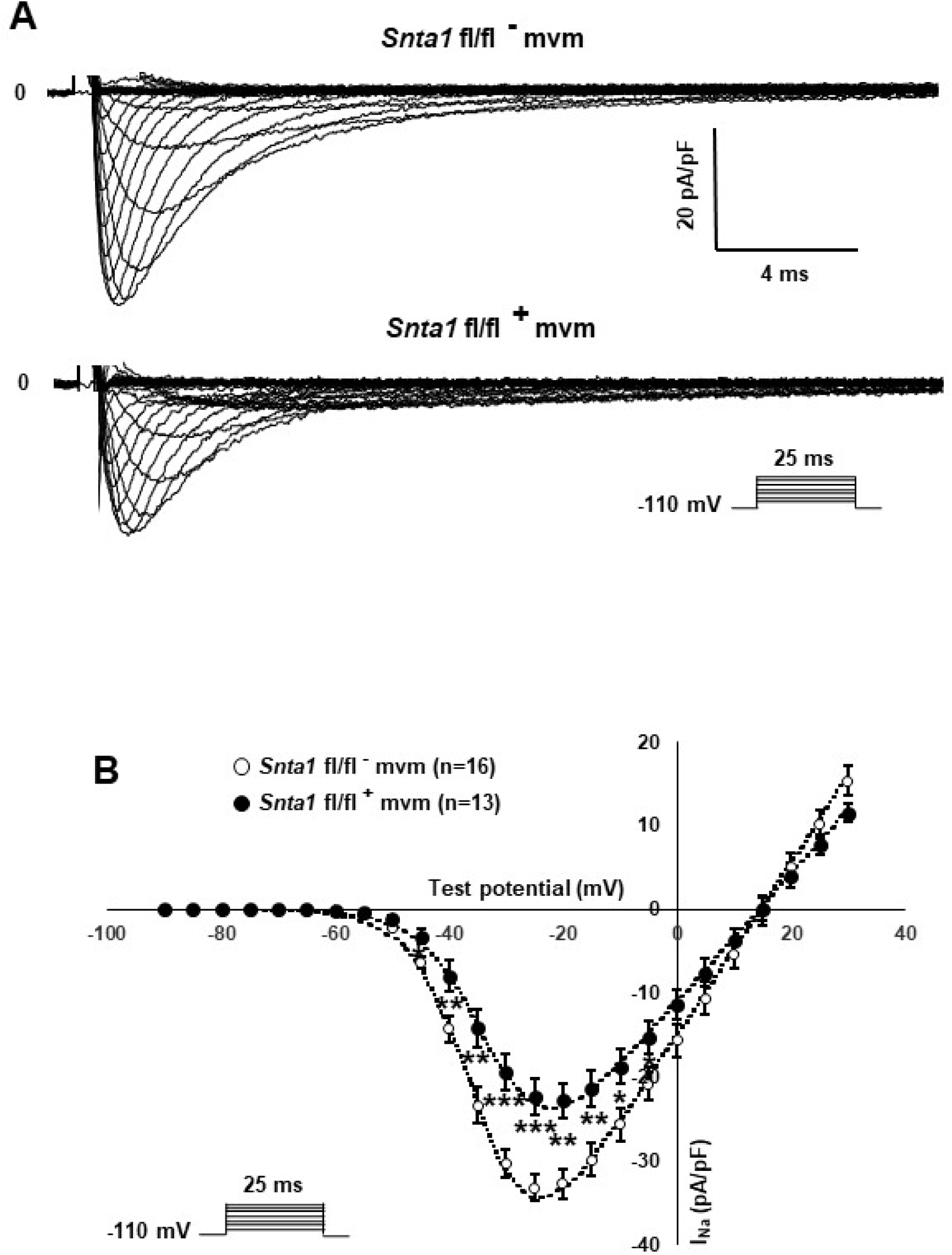
Functional characterization of α1-syntrophin knockdown cardiomyocytes. A, Raw traces of whole-cell sodium currents recorded in adult ventricular cardiomyocytes from wild-type (*Snta1* fl/fl^-^) and α1-syntrophin cardiac specific knockdown cardiomyocytes (*Snta1* fl/fl^+^). B, Whole-cell sodium current I-V curves show a significant decrease in the current density in knockdown cardiomyocytes compared to wild-type littermate cells. (*, **, ***; *p* < 0.05, *p* < 0.01, *p* < 0.001). The number of cells is indicated in parentheses.

**Figure 4.**
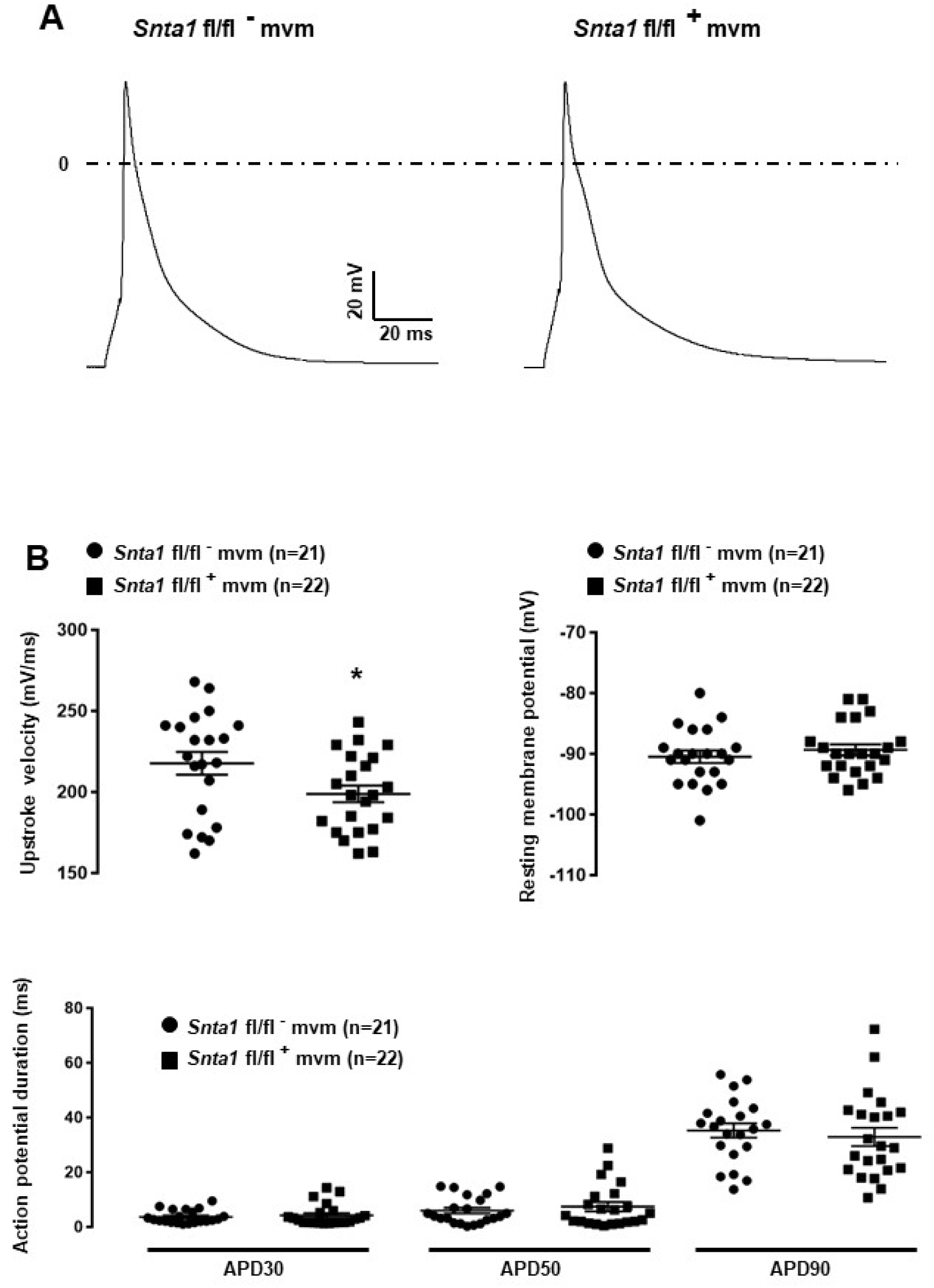
Cardiac action potentials of α1-syntrophin knockdown mice. A, Raw traces of cardiac action potential recorded in adult ventricular cardiomyocytes from wild-type (*Snta1* fl/fl^-^) and α1-syntrophin cardiac specific knockdown cardiomyocytes (*Sntα1* fl/fl^+^). B, Cardiac action potential parameters measured in wild-type (*Snta1* fl/fl^-^)) and α1-syntrophin cardiac specific knockdown cardiomyocytes (*Sntα1* fl/fl^+^) reveal the significant decrease of the maximal upstroke velocity.

**Figure 5.**
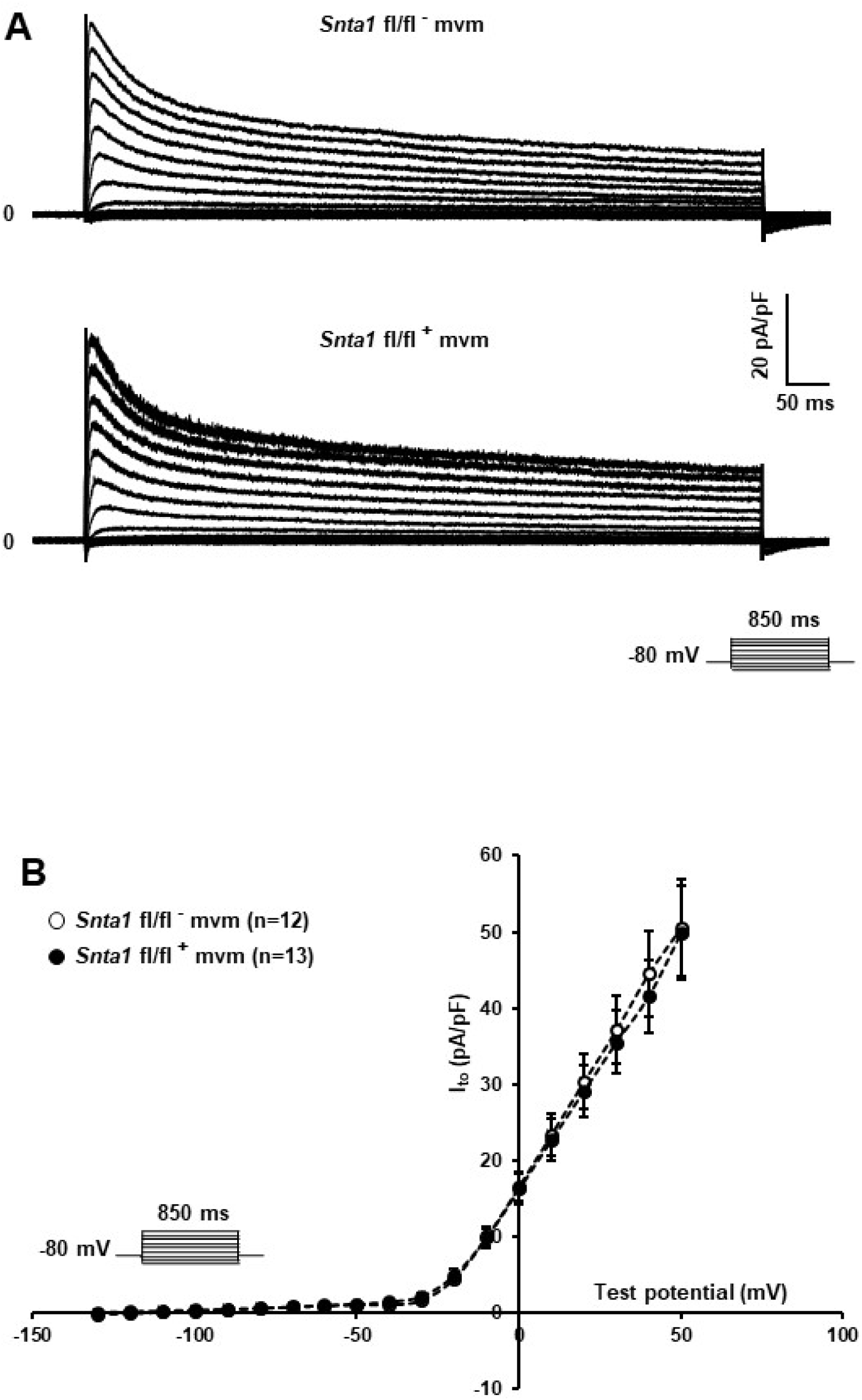
Potassium current I_to_ is not altered in α1-syntrophin knockdown cardiomyocytes. A and B, Raw traces and potassium current I-V relationships show that knockdown of α1-syntrophin has no effect on these parameters in cardiac cells. The number of cells is indicated in parentheses.

**Figure 6.**
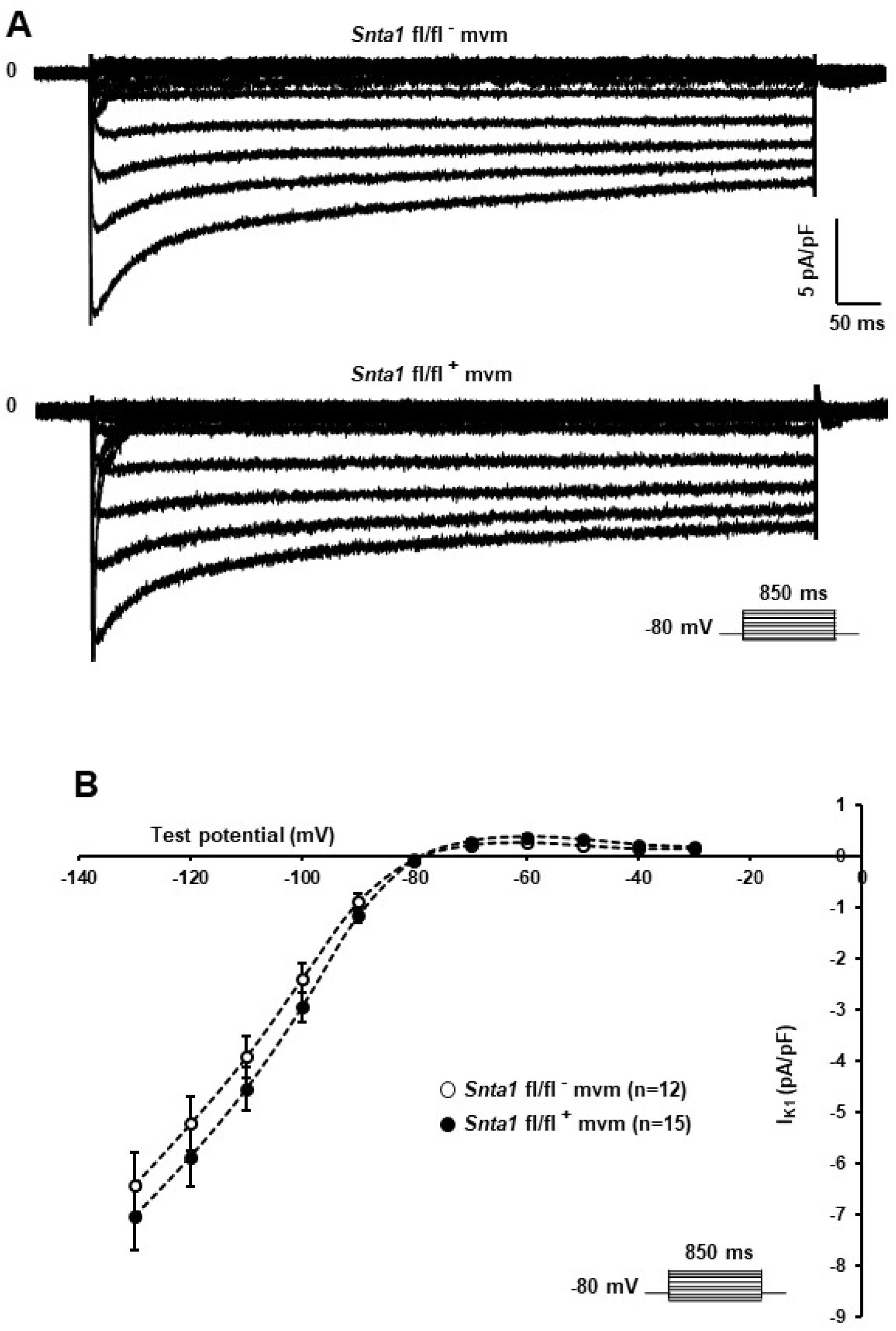
Potassium current I_K1_ is not altered in α1-syntrophin knockdown cardiomyocytes. A and B, Raw traces and potassium current I-V relationships show that knockdown of α1-syntrophin had no effect on these parameters in cardiac cells. The number of cells is indicated in parentheses.

### I_Na_ is reduced but not abolished at the lateral membrane of α 1-syntrophin knockdown cardiomyocytes

Using the cell-attached patch-clamp configuration, we then recorded the I_Na_ specifically at the lateral membrane of α1-syntrophin knockdown cardiomyocytes. As shown in figure 7A and 7B, using an intrapipette solution with 50 nM of TTX to block TTX-sensitive sodium channels (see methods), a sizable TTX-resistant sodium current was recorded in α1-syntrophin knockdown myocytes (Fig. 7A and 7B). This I_Na_ is however smaller (without any alteration of the main biophysical properties) than the currents recorded in control cells (*Snta1* fl/fl^-^ MaxI_Na_: 143 ± 18 pA, n=16; *Snta1* fl/fl^+^ MaxI_Na_: 105 ± 9 pA, n = 22, p < 0.05) (Fig. 7A and 7B and supplemental Fig. 3). Altogether, these results suggest that the reduction of α1-syntrophin expression in ventricular cardiomyocytes leads to a decrease of lateral membrane Na_v_1.5 currents similar to the findings reported by Shy *et al*. in ΔSIV mice [11].

**Figure 7.**
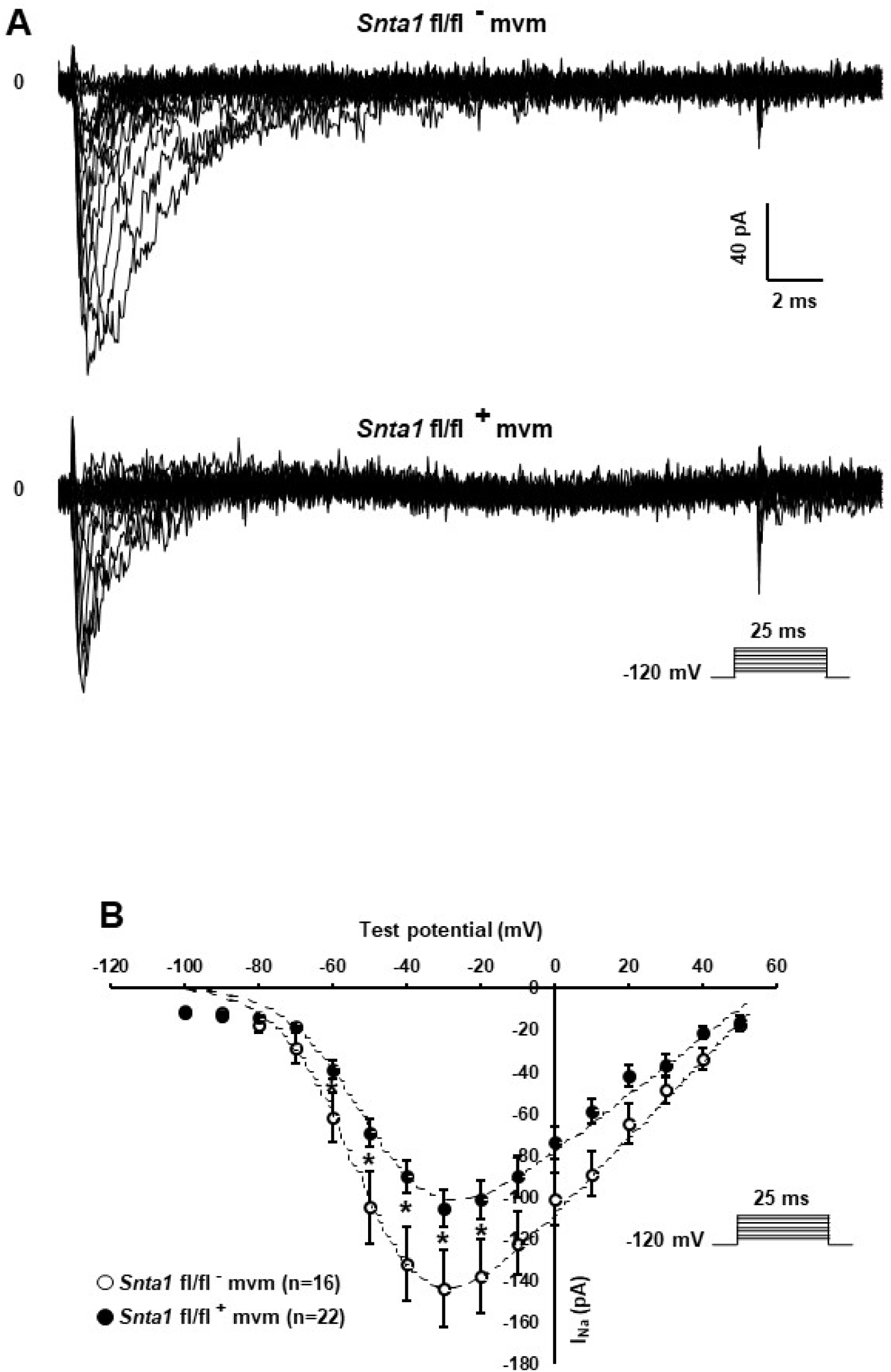
Lateral sodium currents in cell-attached configuration from α1-syntrophin knockdown cardiomyocytes. A, Raw traces of cell-attached sodium currents recorded from adult ventricular cardiomyocytes from wild-type (*Snta1* fl/fl^-^) and α1-syntrophin cardiac specific knockdown cardiomyocytes (*Snta1* fl/fl^+^). B, Sodium current I-V curves from α1-syntrophin cardiac specific knockdown cardiomyocytes (*Snta1* fl/fl^+^) (•) reveal a decrease of the lateral sodium current compared to the control (*Snta1* fl/fl^-^) (○).

### Comparing I_Na_ from α 1-syntrophin knockdown and ΔSIV cardiomyocytes

We then performed I_Na_ recordings using the cell-attached patch-clamp configuration on isolated ventricular cardiomyocytes from wild-type and ΔSIV mice. Again, similar to the observations by Shy *et al*. [11], the TTX-resistant maximum I_Na_ at the lateral membrane was significantly decreased in ΔSIV cardiomyocytes compared to controls (wild-type I_Na_: 198 ± 39 pA, n = 10; ΔSIV I_Na_: 107 ± 14 pA, n = 13, p < 0.05) (Supplemental Fig. 4). As already reported, the biophysical properties (steady-state inactivation and activation curves) did not differ between wild-type and ΔSIV I_Na_ (Supplemental Fig. 5) [11]. Of note, the peak I_Na_ also did not differ significantly between lateral membranes of ΔSIV and α1-syntrophin knockdown cardiomyocytes (ΔSIV MaxI_Na_: 100 ± 8 pA, n = 42; *Snta1* fl/fl^+^ MaxI_Na_: 116 ± 12 pA, n = 26, p > 0.05) (Fig. 8A). In summary, these results suggest that a sizable fraction of the TTX-resistant I_Na_ at the lateral membrane of ven-tricular cardiomyocytes depends neither on the Na_v_1.5 SIV motif nor on α1-syntrophin.

**Figure 8.**
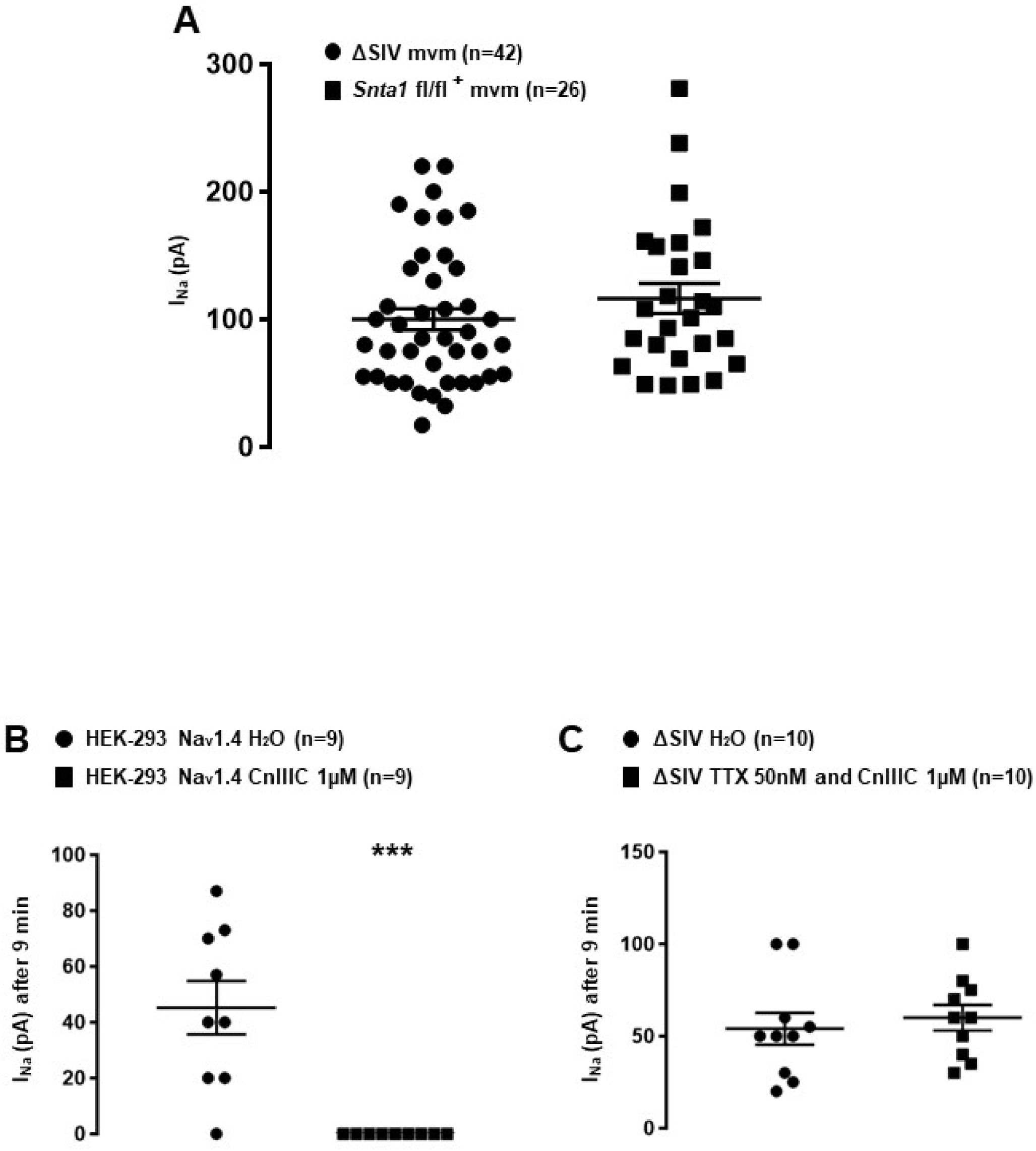
Pharmacological characterization of the remaining sodium current at the lateral membrane of cardiomyocytes. A, Plot chart showing the similarity in average sodium currents recorded at the lateral membrane of cardiomyocytes from ΔSIV and α1-syntrophin cardiac specific knockdown (*Snta1* fl/fl^+^) cardiomyocytes using cell-attached configuration. B, µ-conotoxin CnIIIC (CnIIIC) mediated the reduction in Na_v_1.4 I_Na_ after 9 minutes of application in HEK-293 cell line. C, Tetrodotoxin (TTX) and µ-conotoxin CnIIIC (CnIIIC) did not mediate any effect on I_Na_ recorded on the lateral membrane of ΔSIV cardiomyocytes using the cell-attached configuration (*, **, ***; *p* < 0.05, *p* < 0.01, *p* < 0.001). The number of cells is indicated in parentheses.

### Na_v_ 1.5 channels likely mediate the remaining I_Na_ at the lateral membrane

Several studies report that other Na_v_1.x isoforms may be expressed in cardiac cells in addition to Na_v_1.5, such as Na_v_1.1 [15], Na_v_1.2 [19], Na_v_1.3 [15], Na_v_1.4 [15], Na_v_1.6 [15], and Na_v_1.8 [20]. Although we performed the aforementioned I_Na_ recordings in the presence of 50 nM TTX, which efficiently inhibits Na_v_1.1 (IC_50_ ∼10 nM), Na_v_1.2 (IC_50_ ∼10 nM), Na_v_1.3 (IC_50_ ∼10 nM), and Na_v_1.6 (IC_50_ ∼5 nM) channels, I_Na_ from Na_v_1.4 (IC_50_ ∼25 nM), and Na_v_1.8 (IC_50_ >50 µM) isoforms could still be present. As shown in supplemental figure 4, in recordings from ΔSIV cells, the current-voltage (I-V) relationship presents the maximum I_Na_ at a membrane voltage of around −30 mV. This suggests that the remaining current is not conducted by Na_v_1.8, which maximum current lies at more depolarized values (around +10 mV) [21]. We then addressed the question whether Na_v_1.4 contributes to the remaining current. First, to do so, HEK-293 cells were transfected with a Na_v_1.4 construct, and currents were recorded using the cell-attached patch-clamp approach in the presence of several drugs. As shown in supplemental figure 6A, I_Na_ was only recorded in cells transfected with Na_v_1.4 (Supplemental Fig. 6A). Addition of 50 nM TTX in the intrapipette solution strongly decreased the Na_v_1.4 I_Na_ after three minutes of pipette seal (H_2_O I_Na_: 54 ± 21 pA, n = 7; TTX 50 nM I_Na_: 6 ± 3 pA, n = 9, p < 0.05) (Supplemental Fig. 6B). In parallel, nine minutes after pipette seal, an intrapipette solution containing 1 µM µ-conotoxin CnIIIC, a specific blocker of Na_v_1.4 (IC_50_ ∼1.4 nM) compared to Na_v_1.5 (IC_50_ > 10 µM) [22], completely abolished Na_v_1.4 currents (H_2_O I_Na_: 45 ± 10 pA, n = 9; CnIIIC 1 µM I_Na_: 0 ± 0 pA, n = 9, p < 0.05) (Fig. 8B). Based on these observations, an intrapipette (cocktail) solution containing 50 nM of TTX and 1 µM µ-conotoxin CnIIIC was then used to repeat the I_Na_ recordings from the lateral membrane of ΔSIV cardiomyocytes. As shown in figure 8C, nine minutes after sealing the pipette, no alteration of the I_Na_ amplitude was observed (H_2_O I_Na_: 54 ± 9 pA, n = 10; cocktail I_Na_: 60 ± 10 pA, n = 10, p>0.05) (Fig. 8C). These results suggest that the remaining lateral membrane current in ΔSIV cardiomyocytes is mainly conducted by Na_v_1.5 channels.

### Roles of T-tubular domain in generating the lateral membrane I_Na_

We then aimed to investigate whether lateral membrane I_Na_ is of T-tubular origin, following a protocol described by Brette and colleagues on detubulated cardiac ventricular cells [18]. As shown in figure 9A, di-8-ANEPPS stainings on living non-detubulated wild-This striated pattern was absent in detubulated wildtype cells, validating the detubulation procedure (Fig. 9A). To assess the functional consequences of detubulation, we first recorded the electrical capacitance of the cells, which approximates the surface of the plasma membrane, and, as a positive control, Ca_v_1.2-mediated currents using Ba^2+^ as charge carrier (Fig. 9B, 9C, and 9D). Detubulation significantly decreased the capacitance of the cardiac cells by 30 ± 4%, (non-detubulated: 159 ± 14 pF, n = 7; detubulated: 111 ± 6 pF, n = 7, p < 0.05) (Fig. 9B), and the maximum Ba^2+^ current by 57 ± 7% (non-detubulated I_Ba_: 4377 ± 270 pA, n = 7; detubulated I_Ba_: 1879 ± 306 pA, n = 7, p < 0.05) (Fig. 9C). Consequently, the Ba^2+^ current density (pA/pF) was also significantly decreased by 42 ± 9 % (non-detubulated I_Ba_: 29 ± 3 pA/pF, n = 7; detubulated I_Ba_: 17 ± 3 pA/pF, n = 7, p < 0.05) (Fig. 9D). These results validate the procedure of cardiomyocyte detubulation by formamide treatment. Then, lateral I_Na_ was recorded in non-detubulated and detubulated ΔSIV cardiomyocytes. As shown in figure 10A and 10B, the maximum I_Na_ of detubulated cardiac cells compared to non-detubulated cardiomyocytes was not modified (non-detubulated I_Na_: 95 ± 10 pA, n = 29; detubulated I_Na_: 80 ± 6 pA, n = 29, p > 0.05) (Fig. 10A and 10B). These findings suggest that the remaining lateral membrane I_Na_ in ΔSIV myocytes is not conducted by Na_v_1.5 channels in the T-tubular system, but rather by a distinct lateral membrane pool of Na_v_1.5 channels that is independent of the PDZ-binding motif SIV.

**Figure 9.**
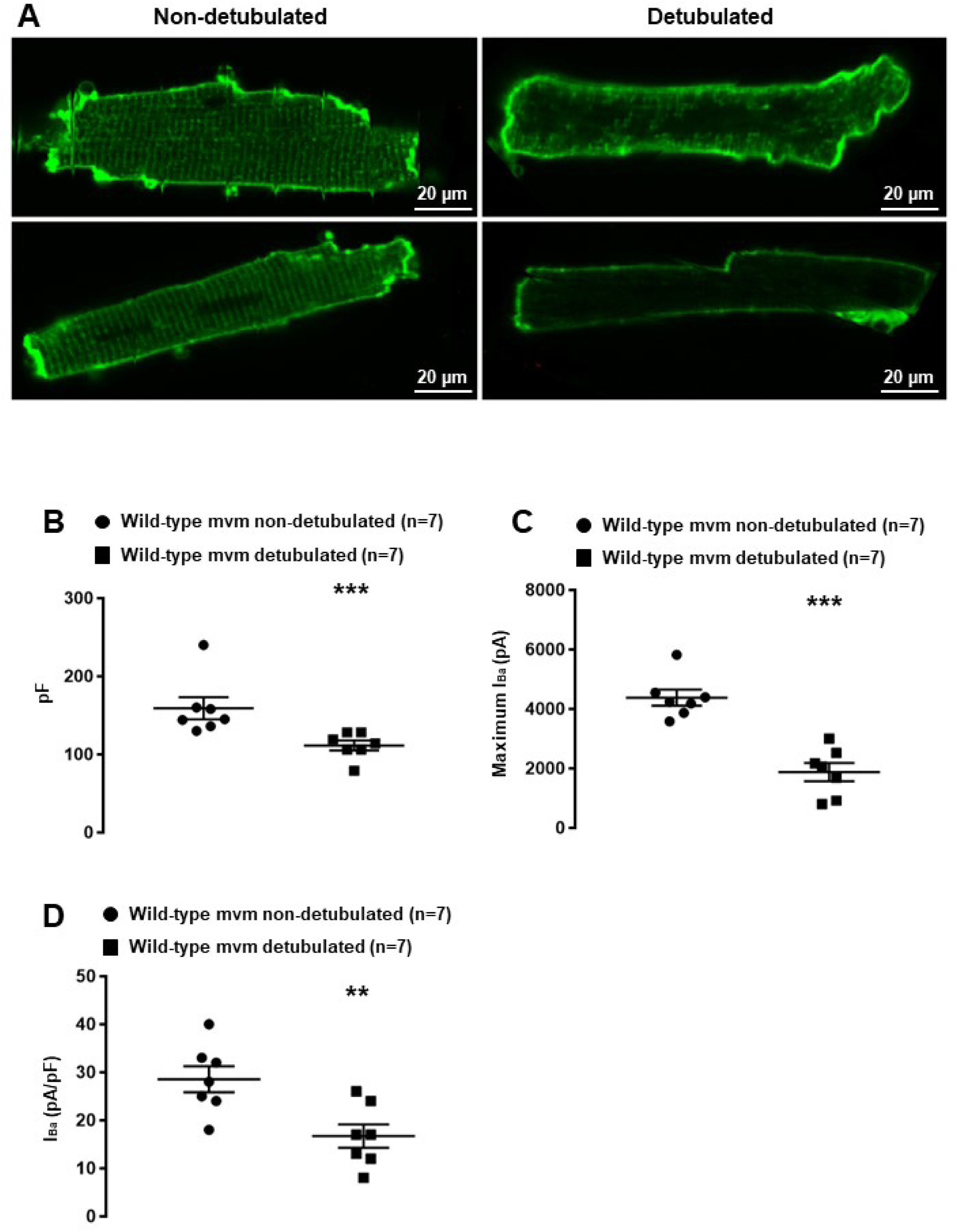
Validation of the detubulation procedure. A, Confocal images showing detubulation mediated by formamide treatment on living cardiomyocytes. B, Effect of detubulation on cell capacitance (pF). C and D, functional consequences of detubulation on barium currents (I_Ba_) conducted by calcium channels. (**, ***; *p* < 0.01, *p* < 0.001). The number of cells is indicated in parentheses.

**Figure 10.**
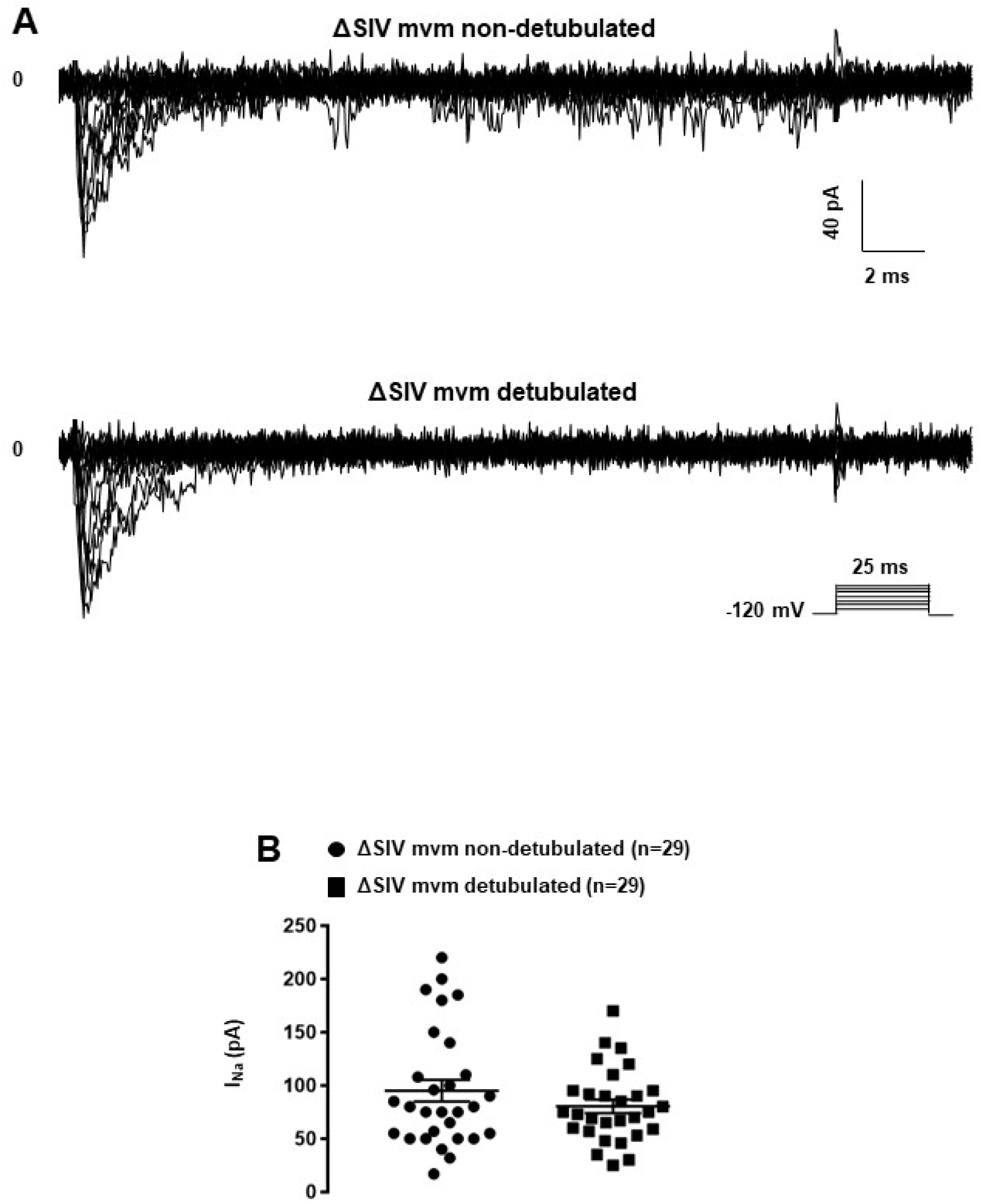
Effects of the detubulation on lateral sodium current. A, Raw traces of cell-attached sodium currents recorded from non-detubulated and detubulated ΔSIV adult ventricular cardiomyocytes. B, Consequences on peak current amplitudes of detubulation on lateral sodium currents from ΔSIV cardiomyocytes. The number of cells is indicated in parentheses.

### Assessing the T-tubular pool of ion channels using the cell-attached patch clamp configuration

Lastly, we assessed whether the cell-attached patchclamp configuration allows the recording of T-tubular currents. In theory, the pipettes used in this study had an opening large enough (∼20 µm^2^) to catch the mouth of 3-5 T-tubules. We recorded I_Ca_ from T-tubules on wild-type ventricular myocytes using an intrapipette solution with only calcium as the charge carrier, since most of the cardiac voltage-gated calcium channels Ca_v_1.2 are located in this membrane domain [23]. Contrary to the expected macroscopic I_Ca_, only single-channel current activities were observed, most likely arising from calcium channels (Fig. 11A and 11B). To characterize these currents, 3 µM of the calcium channel opener Bay-K-8644 was added to the intrapipette solution. Bay-K-8644 significantly increased the number of events (vehicle: 39,677 events, n = 3; Bay-K-8644: 83,697 events, n = 3, p < 0.05) (Fig. 11A and 11B). Altogether, these results demonstrate that the cell-attached patch-clamp configuration on the lateral membrane cannot be used to investigate the function of ion channels in the T-tubular system.

**Figure 11.**
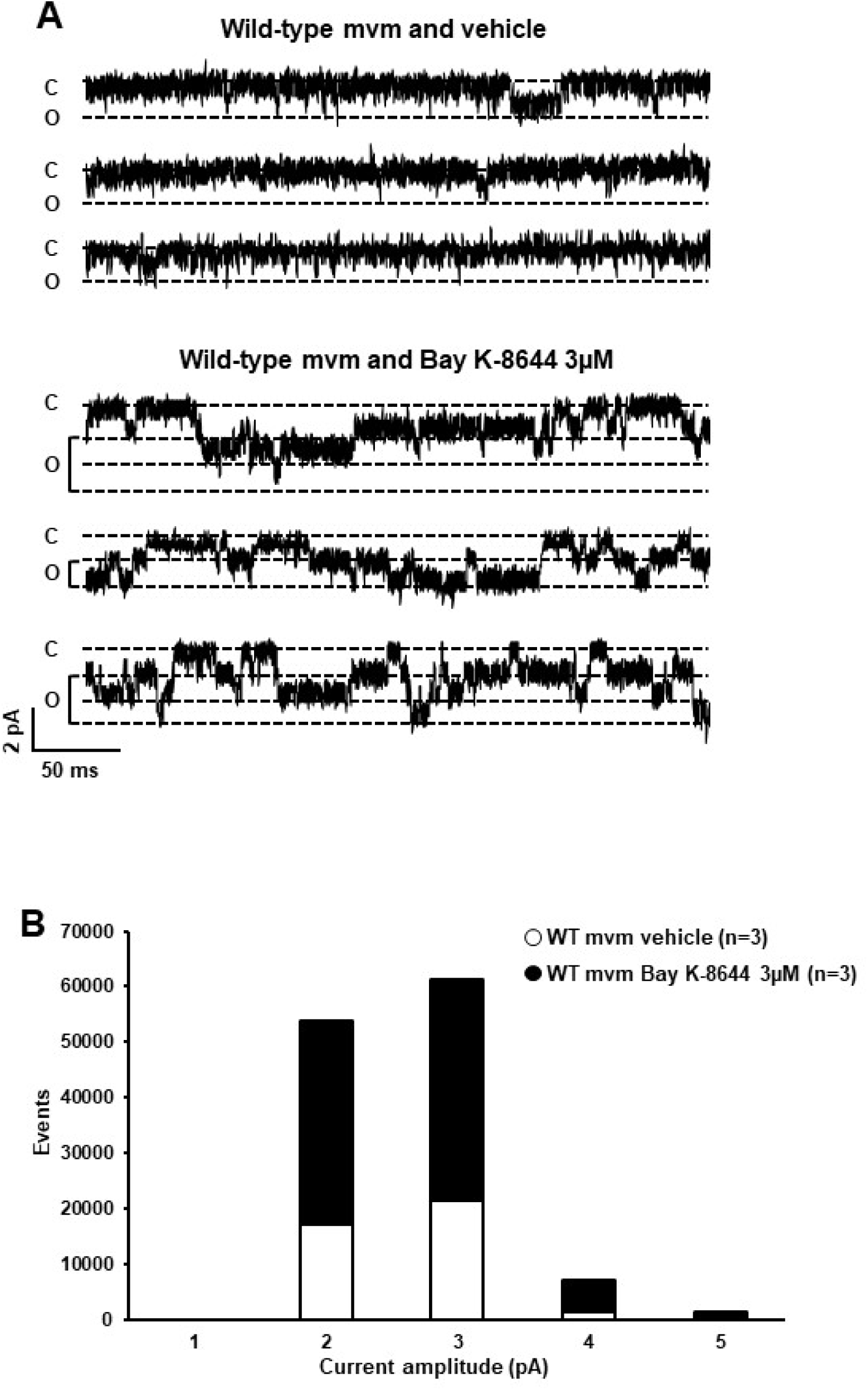
Calcium current recordings and cell-attached configuration. A, Single currents recorded using cell-attached configuration on the lateral membrane of wild-type cardiomyocytes with only calcium inside the pipette. The calcium channel opener Bay K-8644 affects calcium currents, suggesting that this single current is a calcium current. B, Quantification of events observed in A without (○) or with (•) Bay K-8644. The number of cells is indicated in parentheses.

## Discussion

The present study shows that (1) a distinct population of Na_v_1.5 channels at the lateral membrane of cardiomyocyte does not depend on either the PDZ-binding SIV motif or the α1-syntrophin protein; (2) the remaining I_Na_ at the lateral membrane of Na_v_1.5 ΔSIV myocytes is conducted by sodium channels that are most likely not located in the T-tubules; (3) the identity of this sub-pool of sodium channels remaining at the lateral membrane is probably Na_v_1.5; and (4) the cell-attached patchclamp configuration does not allow the investigation of ion channels expressed in the T-tubular system.

Although other Na_v_1.x channels (so-called neuronal isoforms) have been proposed to be expressed in the T-tubular system, recent quantification of RNA transcripts extracted from freshly isolated murine ventricular myocytes suggests that only transcripts encoding Na_v_1.5 and Na_v_1.4 channels are significantly expressed [4]. In ventricular cardiomyocytes, the punctuated pattern observed in stainings by Shy et al. and others might be explained by the presence of Na_v_1.5 inside the T-tubules or at the Z-line level, a structure close to the T-tubules [11, 24]. The presence of a Na_v_1.5 channel population in T-tubules of cardiomyocytes is however still an open question. In the present study, we obtained evidence that suggests that the remaining I_Na_ at the lateral membrane of ΔSIV cardiomyocytes, in which the interaction of Na_v_1.5 with the dystrophin-syntrophin complex is impaired [11], is conducted by an extra-T-tubular compartment. In addition, we concluded that cell-attached patch-clamp experiments at the lateral membrane cannot provide any functional information concerning ion channels in the T-tubular system. Using the cell-attached patch-clamp configuration of the lateral membrane of myocytes, only unitary calcium conductance could be recorded. This observation probably reflects Ca_v_1.2 channels around the opening of T-tubule in the groove of the lateral membrane, as reported by the Gorelik group [25].

Overall, these results confirm the presence of a sub-pool of Na_v_1.5 channels that are not part of the dystrophin-syntrophin complex at the lateral membrane of murine ventricular cardiac cells as suggested by Shy et al. [11]. The novel findings from the cardiac-specific α1-syntrophin knockdown mouse line suggest that the remaining I_Na_ is indeed independent from α1-syntrophin. However, further experiments using a double transgenic mouse model ΔSIV-α1-syntrophin knock down will have to be carried out to indubitably show whether the remaining I_Na_ at the lateral membrane is independent of both the SIV motif and α1-syntrophin.

The remaining I_Na_ in ΔSIV cells recorded in this study, similar to Shy *et al*. [11], represents ∼50% of the total lateral membrane current of mouse cardiomyocytes. Knowing that α1-syntrophin accounts for ∼90% of all syntrophin isoforms expressed in murine cardiac cells, it appears unlikely that the 50% of remaining sodium channels at the lateral membrane interacts only with ∼10% of the other syntrophin isoforms expressed [4]. Altogether, these data suggest the presence of Na_v_1.5 channels independent of SIV motif and α1-syntrophin at the lateral membrane of cardiomyocytes. When interpreting these data, however, we have to keep in mind that the compartmentalization of sodium channels in cardiomyocytes may be species-dependent. Using a similar approach as in the present study, Verkerk and colleagues showed that TTX-resistant voltage-gated sodium channels including Na_v_1.5 are absent at the lateral membrane of rabbit ventricular cardiomyocytes [26]. Moreover, the fact that only I_Na_ and not Ito and I_K1_ was altered in α1-syntrophin knockdown cardiomyocytes is at odds with the recent study by Matamoros *et al*. [13, 17]. This discrepancy may reflect fundamental differences when studying ion channel regulation using *in vitro* vs. *in vivo* models as already observed with the regulation of Na_v_1.5 by SAP97 [6, 8, 9, 17].

The main limitations of the present work are primarily related to the methodological approaches. Future experiments could investigate the precise localization of these syntrophin-independent sub-pools of Na_v_1.5 channels at the lateral membrane with scanning ion conductance microscopy (SICM). In addition, the different techniques used in this study were insufficient to unequivocally demonstrate whether Na_v_1.5 channels are present or absent inside T-tubules. Other techniques should be developed to answer this question. In summary, these results confirm the presence of a sub-pool of Na_v_1.5 channels at the lateral membrane of murine cardiomyocytes, outside the T-tubules, that is independent of α1-syntrophin and the SIV motif of Na_v_1.5. Future experiments will help to understand the functional roles of these distinct pools of sodium channels in specific membrane compartments of cardiomyocytes.

### Source of funding

The research leading to these study results received funding from the European Community’s Seventh Framework Program FP7/2007–2013 under grant agreement No. HEALTH-F2-2009-241526, from EUTrigTreat, and the Swiss National Science Foundation (310030_165741).

## Acknowledgements

We thank Dr. Pascal Peschard, Dr. Lijo Ozhathil, and Sarah Vermij for their useful comments on previous versions of this manuscript. We acknowledge the Microscopy Imaging Center of the University of Bern (MIC) for their contributions to this study.

## Author contributions

Conception and design of the experiments: JSR, and HA. Collection, analysis and interpretation of data: JSR, HA, ME, LG, and SG.

Drafting the article and revising it critically for important intellectual content: JSR, and HA.

---

**Figure S1.**
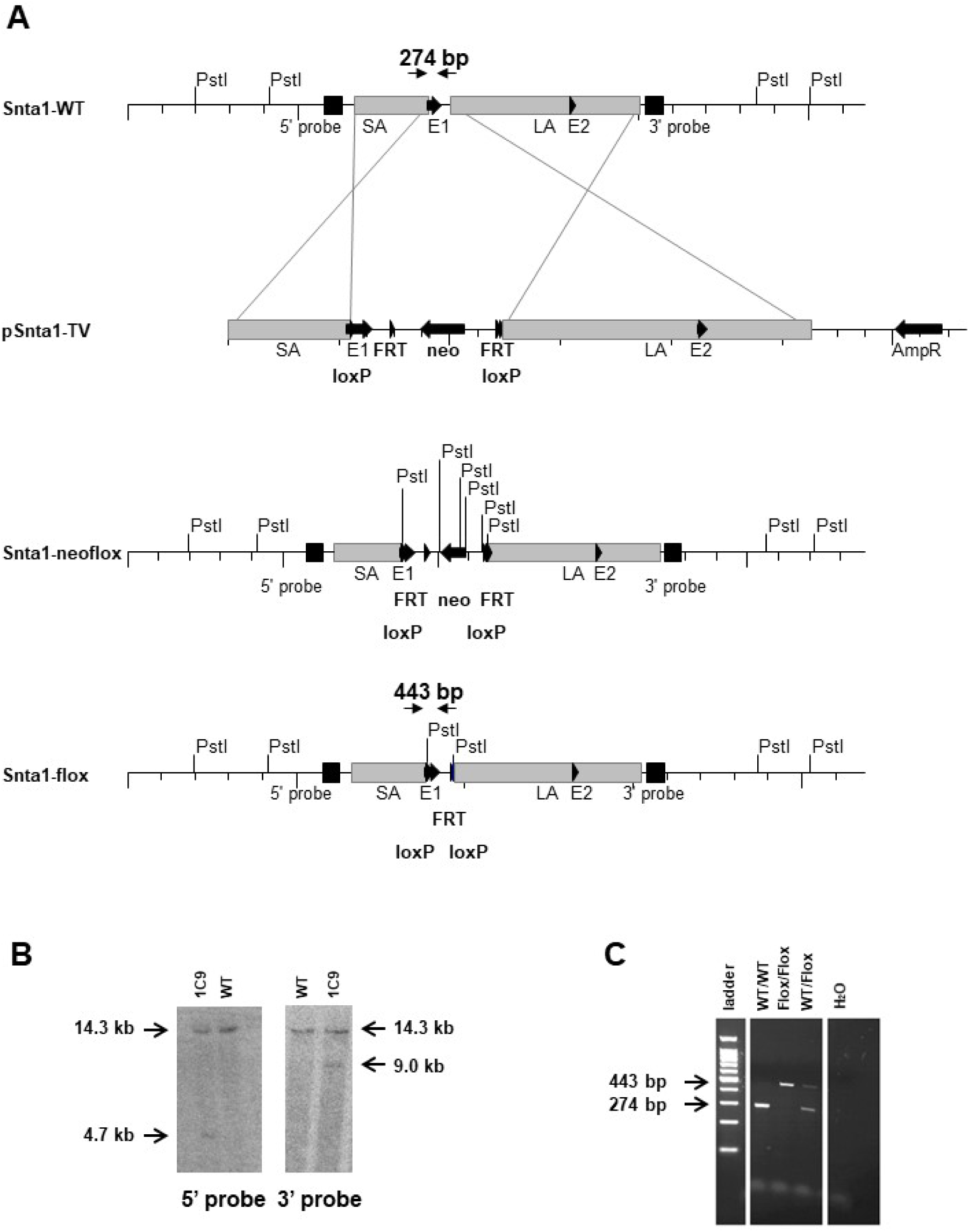
Supplemental Figure 1: Generation of the *Snta1*-flox mouse model. A, The diagram shows homologous recombination of the *Snta1* locus (Snta1-WT) with the targeting vector pSnta1-TV and the resulting targeted Snta1 alleles. Two loxP sites are inserted into the Snta1 locus, flanking a region of 0.7 kb that contains part of exon 1 of Snta1. One loxP site is inserted into the 5′ untranslated region (5′ UTR) of exon 1, whereas the second site is located in intron 1. Directly upstream of the 3′ loxP site, a neomycin resistance cassette (neo) is inserted, which is flanked by FRT-sites, the recognition sites for the Flp-recombinase (Snta1-neoflox allele). Upon breeding to Flp-deleter mice, the neomycin resistance cassette is deleted, and only a single FRT site is remaining in the *Snta1*-flox allele. The short (SA) and long homology regions (LA) are indicated as grey boxes. Positions of primers and sizes of products for genotyping PCR, as well as restriction sites and the probes for Southern blot analyses, are indicated. B, Representative result of Southern blot analysis with genomic DNA from Snta1 wt/wt and Snta1 neoflox/wt (1C9) ES cells cut with PstI and probed using either a sequence upstream of exon 1 or downstream of exon 2 of Snta1. This resulted in a 14.3 kb signal for the wild-type allele with both probes and either a 4.7 kb signal for the 5′ probe and a 9.0 kb signal for the 3′ probe in the Snta1-neoflox allele. C, PCR analysis of all three genotypes using the primer depicted in A. The wild-type (WT) allele resulted in a PCR product of 274 bp, whereas a 443 bp fragment was generated for the Snta1-flox allele (Flox).

**Figure S2.**
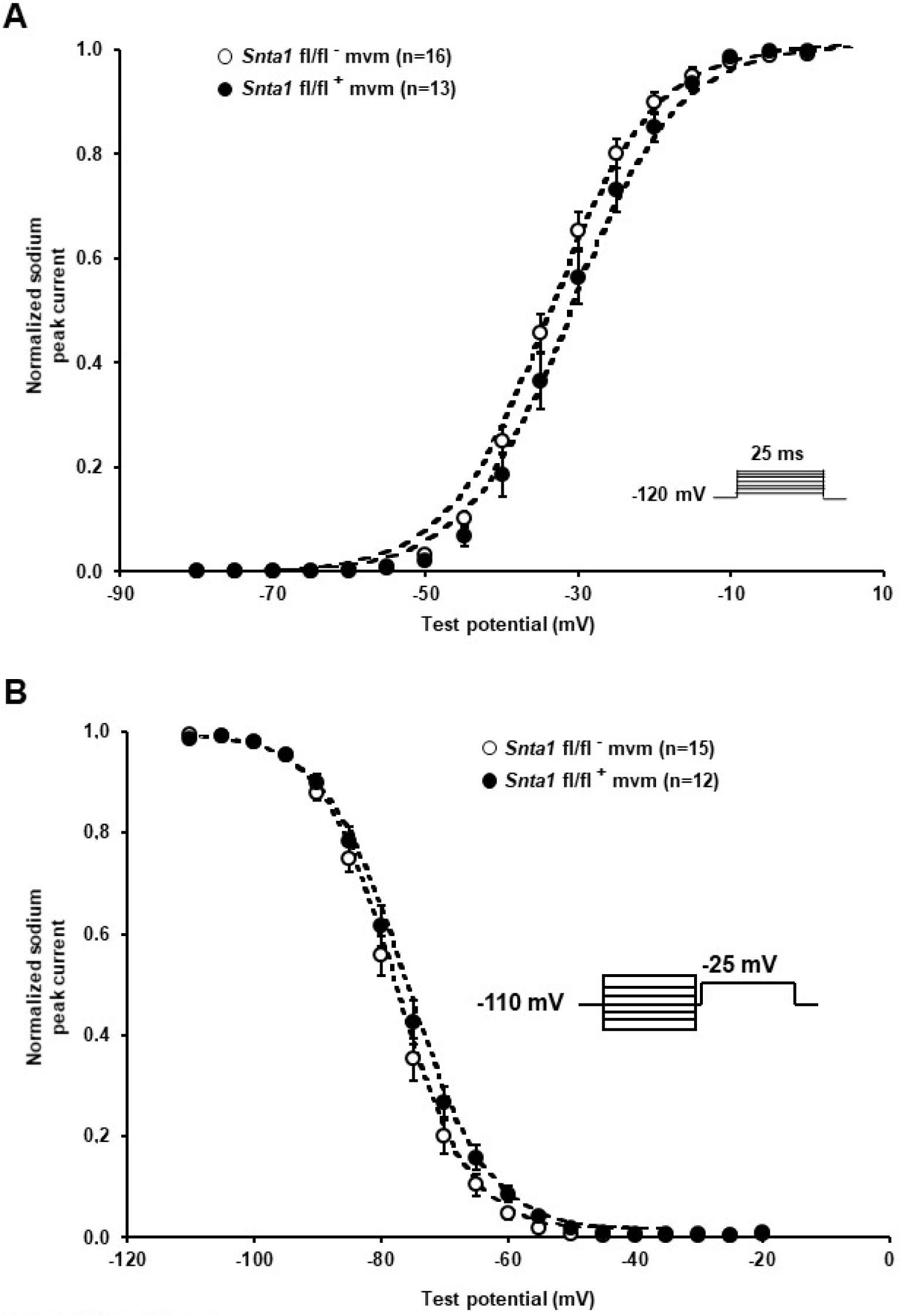
Supplemental Figure 2: A and B, Activation (A) and steady-state inactivation curves (B) calculated from whole-cell recordings of sodium currents show no alteration of the biophysical properties between adult ventricular cardiomyocytes from wild type (*Snta1* fl/fl^-^) and α1-syntrophin cardiac specific knockdown cardiomyocytes (*Snta1* fl/fl^+^).The number of cells is indicated in parentheses.

**Figure S3.**
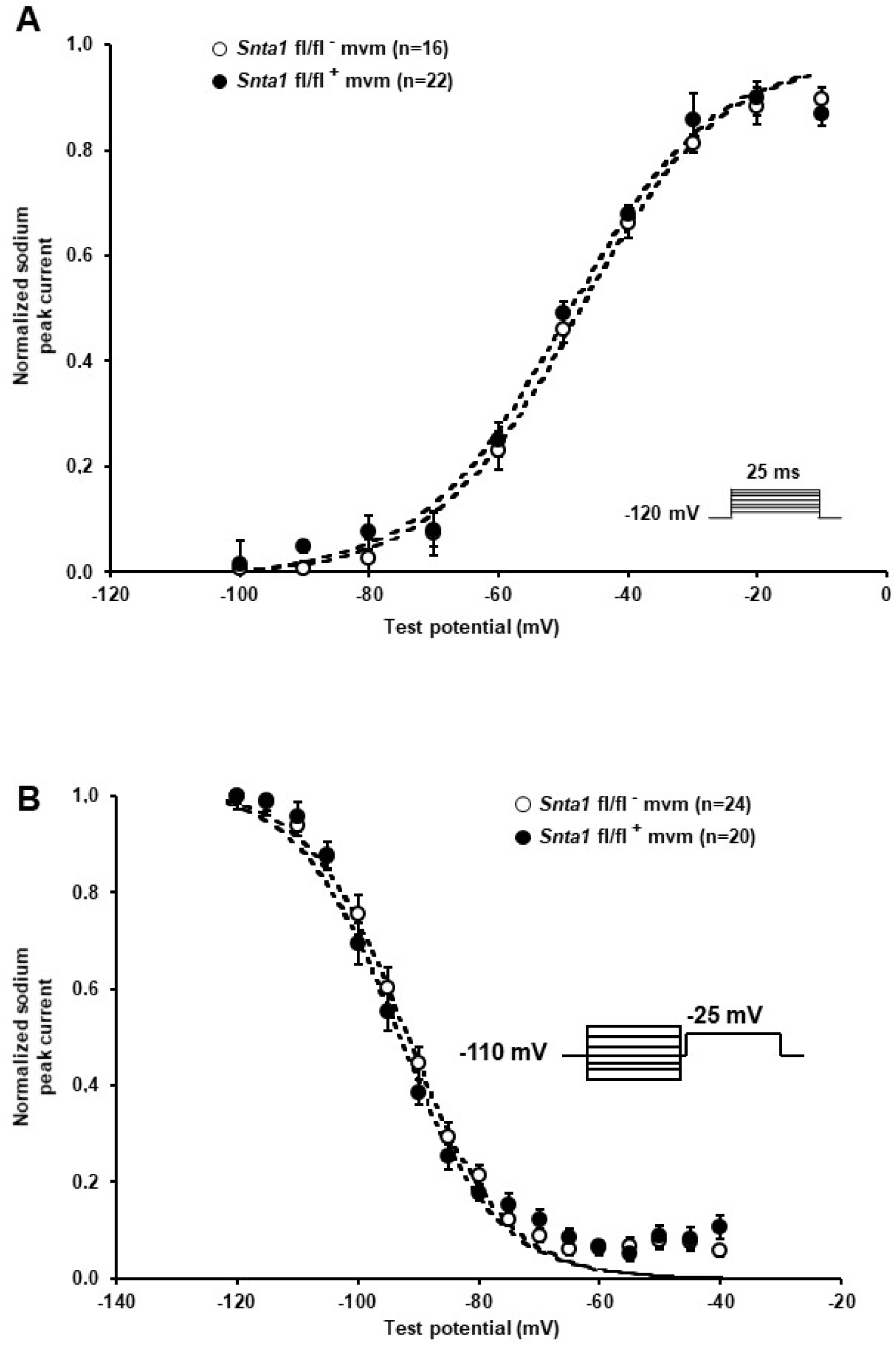
Supplemental Figure 3: A and B, Activation (A) and steady-state inactivation curves (B) calculated from cell-attached recordings of sodium currents show no alteration of the biophysical properties between adult ventricular cardiomyocytes from wild-type (*nta1* fl/fl^-^) and α1-syntrophin cardiac specific knockdown cardiomyocytes (*nta1* fl/fl^+^). The number of cells is indicated in parentheses.

**Figure S4.**
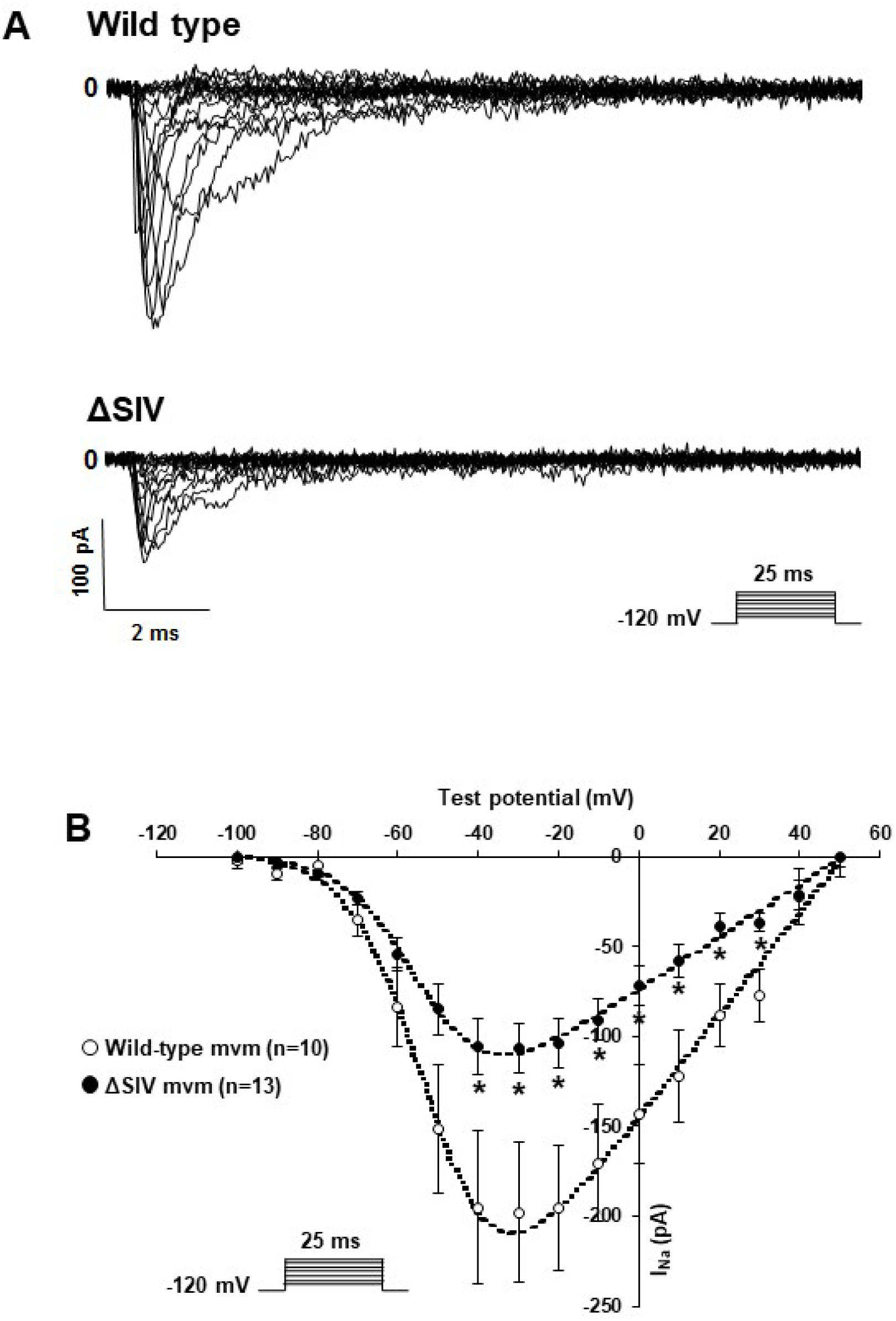
Supplemental Figure 4:Lateral sodium current in ΔSIV cardiomyocytes. A, representative cell-attached recordings of sodium currents from the lateral membrane of wild-type and ΔSIV cardiomyocytes. B, Current-voltage (I-V) curve showing a significant decrease in the sodium current recorded in ΔSIV cardiomyocytes (•) compared to the control (○) (*; *p* < 0.05). The number of cells is indicated in parentheses.

**Figure S5.**
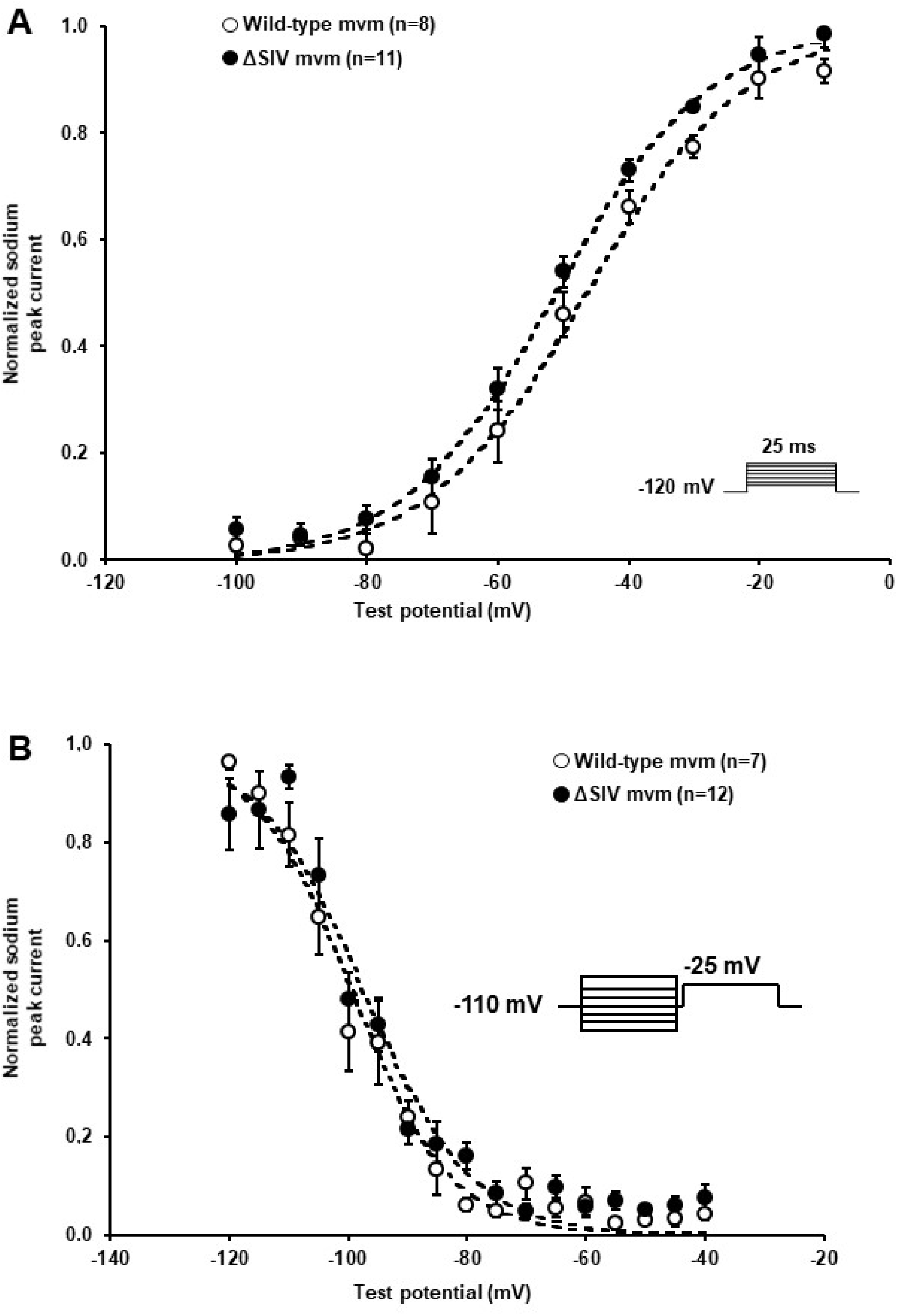
Supplemental Figure 5: A and B, Activation (A) and steady-state inactivation curves (B) calculated from cell-attached recordings of sodium currents show no alteration of the biophysical properties between adult ventricular cardiomyocytes from wild-type and ΔSIV cardiomyocytes. The number of cells is indicated in parentheses.

**Figure S6.**
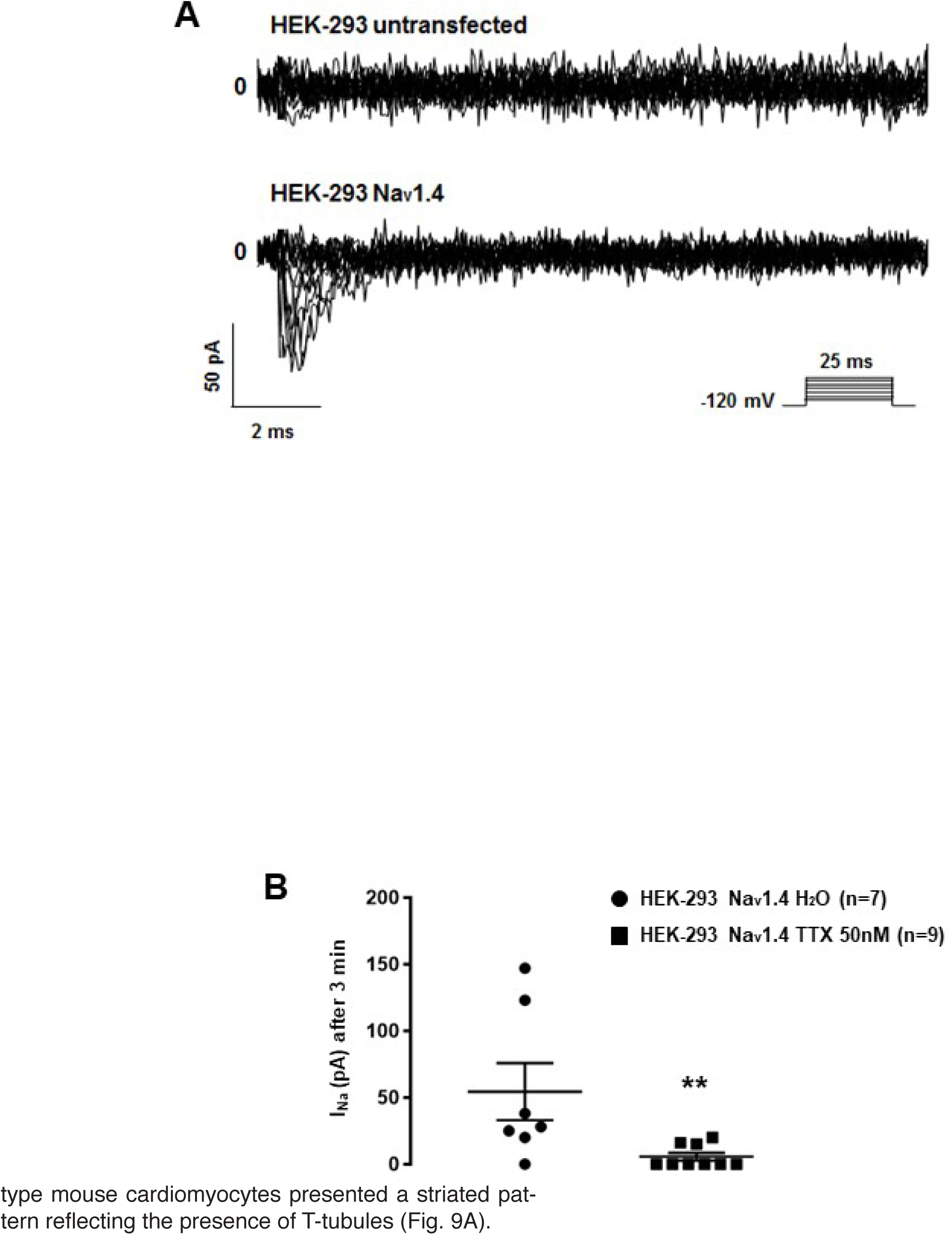
Supplemental Figure 6:Pharmacological block of Na_v_1.4 current. A, Representative cell-attached Na_v_1.4 currents recorded in transiently transfected HEK-293 cells. B, I_Na_ reduction mediated by tetrodotoxin (TTX) after 3 minutes of application (**; *p* < 0.01). The number of cells is indicated in parentheses.

